# The Sorghum Lipid Database (SoLD): population-scale lipidomics linking environmental and genetic variation in the Sorghum Association Panel

**DOI:** 10.64898/2026.06.05.727974

**Authors:** Nirwan Tandukar, Ruthie Locklear, Richard Boyles, Zach Brenton, Katherine Louie, Rubén Rellán-Álvarez

## Abstract

Sorghum (*Sorghum bicolor*) is a climate-resilient crop whose acclimation to nutrient limitation and low temperature likely involves extensive lipidome reconfiguration. Lipids are key membrane components, carbon and energy stores, and mediators of stress signaling, yet population-scale lipidomics data for sorghum are limited. We present the Sorghum Lipid Database (SoLD), a curated lipidomics resource from the Sorghum Association Panel grown under two field regimes, (i) a nutrient-sufficient with usual planting date environment (control) and (ii) a low input treatment with reduced nitrogen and phosphorus, earlier planting, and no pesticide application (low input). Using high-resolution LC–MS, we quantified 266 lipid species and found the lipidome broadly similar across trials, with four low-input-associated signatures relative to control, (i) depletion of sulfoquinovosyldiacylglycerol, (ii) triacylglycerol enrichment, (iii) phospholipid redistribution centered on phosphatidylserine, and (iv) a shift in the lysophosphatidylcholine-to-lysophosphatidylethanolamine ratio. Genomic heritability was zero at the median under control against 0.171 under low input, and more of the lipidome could be mapped under low input. GWAS of lipid species, class sums, and class ratios recovered loci that were largely trial-specific, with candidate genes assigned by linkage disequilibrium and prioritized by LD-aware ontology enrichment and reaction mapping. Control-associated loci yielded twenty enriched gene sets, all confined to triacylglycerol traits and led by cutin biosynthesis, whereas low-input-associated loci yielded fifty-one across five lipid classes, centered on nutrient transport and signaling, cold-responsive regulators, and jasmonate and oxylipin genes, and including a chromosome 3 block carrying two LOW PHOSPHATE ROOT homologues. SoLD thus connects sorghum lipid diversity to environmental and genetic variation, and lets users move from lipid class to individual species and associated candidate loci. It is freely accessible at https://nirwan.shinyapps.io/SAP-Lipidomics-Database/.

## Introduction

Plants routinely encounter environmental stresses that reduce yield, including nutrient limitation, low temperature, and disease. Nitrogen (N) and phosphorus (P) deficiencies constrain growth and nutrient allocation, while cold disrupts membrane properties and metabolism (Schlüter et al. 2012; Silveira *et al*. 2018; Emendack et al. 2021; Ding et al. 2019; Guo et al. 2025). Lipids are key to these responses as membrane components, energy reserves, and signaling precursors. Stress-induced lipid remodeling affects membrane stability, photosynthesis, storage lipids, and lipid signaling (Bohn et al. 2007; Siebers *et al*. 2015; Hou et al. 2016; Rawat et al. 2021; Li et al. 2020a; Bhattacharya 2022; Xu et al. 2025). Lipids also contribute to plant defense against pathogens, particularly, cuticular and membrane lipids form physical barriers to infection (Kuźniak and Gajewska 2024a). In sorghum, fungal diseases such as anthracnose, grain mold, charcoal rot, downy mildew, and rust can substantially affect yield and grain quality, making disease pressure an important component of field stress (Khaskheli *et al*. 2025). Under P limitation, phospholipids are often replaced by non-P galactolipids and sulfolipids such as DGDG and SQDG, whereas cold can change acyl-chain composition, phospholipid turnover, and triacylglycerol (TG) accumulation (Essigmann *et al*. 1998; Härtel *et al*. 2000; Jouhet *et al*. 2004; Kelly and Dörmann 2002; Tan *et al*. 2018; Pfaff *et al*. 2020; Yan *et al*. 2025; Zhang *et al*. 2020). These shifts are coordinated by nutrient and stress pathways, including SPX–PHR phosphate signaling, nitrate transport (NRT2/NPF), sulfate transport for plastid sulfur supply, and enzymes driving phospholipid turnover and DG–TG cycling (Lv *et al*. 2014; Lupini *et al*. 2016; Sorgonà *et al*. 2011; Xia *et al*. 2015; Chen *et al*. 2019; Bose *et al*. 2021; Ngo *et al*. 2019; Tan *et al*. 2018; Kelly *et al*. 2011; Yen *et al*. 2008). However, crop-scale lipidomics linking lipid remodeling, nutrient stress, biotic stress, and natural variation is still limited, especially in sorghum.

Sorghum is a stress-tolerant cereal with extensive diversity. The Sorghum Association Panel (SAP) captures broad botanical, geographic, and genomic variation for studying environmentally and agronomically important traits (Casa et al. 2008; Boatwright et al. 2022). Because population lipidomics is sensitive to technical and field variation and depends on consistent annotation, we applied SERRF normalization, SpATS spatial correction, and GNPS-assisted annotation to improve reproducibility and interpretation (Fan et al. 2019; Rodriguez-Alvarez et al. 2018; Wang et al. 2016; Molenaar et al. 2019).

Interpreting large candidate gene sets from lipid GWAS is challenging because LD-linked genes are not independent. A single associated region may contain multiple neighboring, similarly annotated genes, inflating enrichment. GWAS-focused tools (INRICH, SNP2GO, Gowinda) mitigate this by accounting for LD, SNP structure, and gene-level biases (Lee et al. 2012; Szkiba et al. 2014; Kofler and Schlötterer 2012). Accordingly, we tested GO enrichment using an LD-aware interval-permutation null and separated terms supported by multiple independent intervals from those driven by single loci.

We present the Sorghum Lipid Database (SoLD), a curated lipidomics and GWAS resource for the SAP. Leaf samples (V6– V8) were profiled in two field trials: a 2019 control (CTL; adequate nutrients, standard planting) and a 2022 low-input regime (LIN; reduced N and P, earlier planting to induce cold-associated stress, no chemical management). Because CTL and LIN were grown in different years, contrasts are treated as between-trial differences, not causal treatment effects. SoLD integrates LC– MS lipidomics, compositional analysis, lipid ontology, GWAS of species- and class-level traits, LD-aware GO enrichment, LINEX reaction mapping, and an interactive Shiny app. This framework identifies four LIN-associated signatures: SQDG depletion, TG enrichment, PS-centered phospholipid redistribution, and altered LPC/LPE balance, and prioritizes candidate functions spanning nutrient acquisition, DG–TG cycling, phospholipid turnover, cold response, and jasmonate signaling. Each part of the LIN regime is matched by candidate loci, with nitrate transporters for lower nitrogen, phosphate-status signaling proteins for lower phosphorus, a cold-induced transcription factor block for the earlier planting, and jasmonate-pathway genes for the absence of chemical treatment, alongside both sides of DG–TG cycling, including *DGAT1*. SoLD supports exploration of sorghum lipid variation, candidate loci, and accessions for functional validation; the app is available at https://nirwan.shinyapps.io/SAP-Lipidomics-Database/.

## Materials and methods

### Plant Material and Growth Conditions

We used the Sorghum Association Panel, a genetically diverse collection of sorghum accessions representing major botanical races and broad phenotypic diversity (Casa *et al*. 2008; Boatwright et al. 2022). The analyses use the accessions with both genotype and phenotype data for GWAS, 351 under CTL and 357 under LIN, with 351 common to both trials. We evaluated SAP accessions across two different field settings during two growing seasons (2019 and 2022) at the Pee Dee Research and Education Center, Clemson University, Florence, South Carolina. The “control” condition, herein denoted as CTL, involved standard agronomic inputs with sufficient levels of nitrogen (N) and phosphorus (P) along with a typical planting schedule. In contrast, the “low input” condition, denoted as LIN, featured reduced N and P, earlier planting to mimic a cold stress environment, and no pesticide application.

### Schematic diagram

The schematic in Supplementary S1 Fig was created using FigureLabs.ai (https://www.figurelabs.ai/).

### Lipidomics Analysis

#### Sample Preparation and Extraction

Samples were prepared following the standard extraction protocols explained in the paper by (Barnes *et al*. 2022).

#### Liquid Chromatography-Mass Spectrometry (LC-MS) Analysis

Lipid extracts were analyzed at the Joint Genome Institute/Lawrence Berkeley National Laboratory using their standard LC-MS/MS lipidomics workflow. The detailed chromatographic conditions, acquisition parameters, instrumentation, and quality-control procedures are given in the published JGI/LBNL protocol (Louie *et al*. 2025) and in the instrument metadata deposited with the raw data (MassIVE MSV000101940).

#### Feature Detection and Data Processing

Raw LC-MS data was processed using MZmine 2 (Pluskal *et al*. 2010). The pipeline encompassed peak detection, chromatogram construction, deconvolution, and isotope filtering, producing a detailed feature table containing mass-to-charge ratio-retention time (mz-rt) pairs. Isotopic peaks were excluded to minimize redundancy, as a single metabolite can yield multiple co-eluting ions, such as adducts and in-source fragments. The remaining mz-rt duplicates were resolved at the annotation stage described below. We recognize that this degeneracy may result in more detected features than the true number of metabolites present, a factor we accounted for during metabolite identification.

#### Blank/extraction-control filtering, intensity thresholds, and sparsity pruning

Features were removed if their maximum intensity in biological samples (*b*) was < 10 the maximum in extraction controls (*a*), unless any biological sample exceeded the control maximum. We also required a minimum mean intensity of ~10^6^ peak height. Samples with ≥70% zero or missing features were excluded before normalization to avoid unstable scaling and spurious differential signals. A lipid was considered detected in a condition if it passed these filters using only that condition’s samples. The lipids passing in both were shared, otherwise condition-specific.

#### MS/MS spectral library matching and cross-referencing of IDs

For each MS/MS feature, the most intense spectrum was searched against GNPS, while unmatched features were removed. GNPS library hits provide putative MSI level 2/3 identifications and may reflect isomers or near-mass analogs, so all species-level annotations are tentative and should be confirmed with authentic standards. If multiple GNPS hits occurred, the top MQScore (cosine similarity) was kept as the primary annotation, while lower-ranked candidates were retained for verification but excluded from downstream statistics.

#### Systematic Error Removal Using Random Forest (SERRF)

Following the cleaning process, the data were then used for SERRF normalization (Fan *et al*. 2019). We used the SERRF server (https://slfan2013.github.io/SERRF-online/) to obtain the normalized output. After applying SERRF, only biological samples were preserved. Any zeros were substituted with two-thirds of the minimum nonzero value for that feature to prevent potential infinite logarithmic transformations.

#### Spatial Correction

We removed spatial trial effects after SERRF with the R package `SpATS` (Rodriguez-Alvarez et al. 2018), which fits a 2D P-spline ANOVA surface over field coordinates (mean plus smooth row, column, and row-by-column terms). We used SpATS-adjusted intensities (residuals + fitted mean) for down-stream analyses, preserving the original scale, avoiding negative values required by the %TIC, CLR, and log-ratio transforms, and correcting positional artifacts (e.g., edge effects).

#### Lipid Quantification

The identified lipid species were assigned to the eight LIPID MAPS categories and to classes and subclasses within them, following the LIPID MAPS classification (Fahy *et al*. 2009) (Supplementary Table S1). Lipid class refers throughout to the abbreviation used in a species name, such as PC or TG. In each sample, the total intensity for a class was obtained by summing the species within it and normalizing it to the sample’s total ion current, giving relative abundances as percentages of TIC, which were then averaged across samples within each condition.

For genetic analyses, class totals were computed from SpATS genotype BLUPs to preserve spatial correction. Because BLUPs are deviations from the panel mean and are negative for 60% of genotype-by-class combinations, we added the class intercept 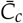 (the panel mean total for class *c* on the input intensity scale) before log transformation (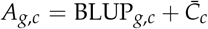). This yielded strictly positive intensities in both trials, so no pseudo-count, shift, or denominator floor was used.

#### Lipid ratio calculation

For each sample *i* and lipid class *c*, species intensities were aggregated to class totals,

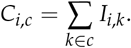

Log-ratios are scale-invariant, so TIC normalization was not needed. Pairwise class log-ratios were

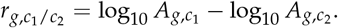

Because 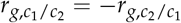, each class pair yields the same trait up to sign and identical association statistics, so we kept only one ordering per pair. With 17 classes, this gives 17 class sums and 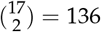 ratios, for 153 traits per trial.

#### Compositional transforms

Class totals and the pairwise class log-ratios defined above are the traits carried forward to GWAS. For species-resolved analyses we used the centered log-ratio transformation, CLR(*x*_*i*_)= ln(*x*_*i*_ /*g*(**x**)), where *g*(**x**) is the geometric mean of species-level abundances in the sample, which removes the constant-sum constraint of compositional data.

### Principal Component Analysis (PCA)

PCA was performed on individual lipid species within each experimental condition. Lipid intensities were first normalized to TIC to obtain relative abundances, then transformed using CLR. The CLR-transformed data were mean-centered and scaled to unit variance, and PCA was conducted in R using the *stats* package. The first two principal components were retained for visualization.

### Genome-wide Association Studies (GWAS)

GWAS analyses were carried out for each lipid trait under each condition using the mixed linear model (MLM) featured in GEMMA (v0.98.5) (Zhou and Stephens 2012). To address population structure and relatedness, a kinship was computed from Sorghum Lipid Database (SoLD): a database curation for lipid information for Sorghum Association Panel SNP genotype data. Genotypes were filtered to biallelic SNPs with a minor allele frequency ≥0.05, and SNPs with more than 80% heterozygous calls were removed. Besides individual traits, GWAS was also applied to summed lipid classes and all possible ratios. The first three genotype principal components were fitted as fixed covariates. A Bonferroni-corrected genome-wide significance threshold was employed to account for multiple comparisons. Based on approximately 6.11 million tested SNPs under CTL and 6.08 million under LIN, this corresponded to *p* ≤8.19 ×10^−9^ for CTL (−log_10_(*p*) ≥8.09) and *p* ≤8.22 ×10^−9^ for LIN (−log_10_(*p*) ≥8.09).

### Gene Annotation

SNPs were aligned to the *Sorghum bicolor* v3.1 (BTx623) reference genome. Because LD in the sorghum association panel spans substantial distances (Boatwright *et al*. 2022), Bonferroni-significant SNPs often tag multiple linked genes. We therefore grouped significant SNPs into association regions at *r*^*2*^≥ 0.4, balancing overly narrow regions at stringent cutoffs against inflated gene counts at permissive ones. Because candidate-gene counts are more sensitive to this threshold than interval counts, all interval-level analyses (GO enrichment and cross-trial overlap) used fixed 250 kb intervals rather than gene IDs. For each candidate gene, we counted linked lipid traits, merging genes that shared a trial, trait layer, chromosome, and best *p*-value into a single signal. These counts summarize the candidate tables but were not used to prioritize genes because lipid phenotypes are correlated by design.

### Botanical race and population-structure testing

Botanical race, working group, and marker-based genetic cluster assignments for the SAP were taken from panel passport data. For race analysis, accessions were grouped into the five classical races (Bicolor, Caudatum, Durra, Guinea, Kafir) plus a Mixed group containing all hyphenated intermediate designations. Accessions lacking a race assignment and the single *S. bicolor* subsp. *verticilliflorum* accession were excluded. The second grouping used the *K* = 6 genetic cluster assignment. Fourteen traits were tested per condition. Groups with fewer than five accessions in a condition were excluded from that test. Differences among groups were assessed with Kruskal–Wallis tests within each condition, and *p*-values were adjusted across the 14 traits per condition using the Benjamini–Hochberg (BH) procedure. Effect sizes were computed as *ε*^2^ = (*H k* + 1)/(*n k*), where *H* is the tie-corrected statistic, *k* the number of groups, and *n* the number of accessions, while negative values were truncated to zero.

The stability of the class composition within a trial was assessed by resampling rather than by hypothesis testing. Neither procedure poses a null hypothesis, computes a test statistic, or yields a *p*-value. The first is a leave-one-group-out sensitivity analysis. Within a trial, every accession belonging to one botanical race group was deleted, and the mean %TIC of each of the 13 lipid classes was recomputed from the remaining accessions once for each of the six groups, giving 78 recomputations per trial. We report the smallest and largest value any omission produced and the largest absolute departure from the full-panel mean. The second is nonparametric subsampling. One thousand draws of 80% of a trial’s accessions were taken without replacement (281 of 351 under CTL, 286 of 357 under LIN; random seed fixed at 1). The same 13 class means were recomputed for each draw, and the 2.5th and 97.5th percentiles of the resulting dis-tribution were calculated as a percentile interval. That interval describes how far the descriptive mean moves under resampling. Both procedures run entirely within a single trial and involve no comparison between trials.

### Candidate-locus definition and cross-trial overlap testing

Cross-trial overlap was calculated from LD-mapped candidates for the individual-lipid layer, the class sum ratio layer, and their union. Gene-level overlap used the 34,027 annotated *S. bicolor* v3.1 genes as background. Because LD mapping assigns all genes in a window to an association and inflates gene counts, we also analyzed overlap at the locus level. The genome was split into non-overlapping 250 kb windows, and a window was a candidate locus for a condition if it contained at least one candidate gene. The locus-level background was defined as windows containing at least one annotated gene (2,279). At both levels, overlap was tested with a one-sided hypergeometric (Fisher right-tail) test and summarized by *p*-value, fold enrichment over the expected overlap |*A*| |*B*| /*N*, and the Jaccard index |*A∩B*| / |*A∪ B*|.

For per-class analyses, each candidate gene was assigned lipid classes based on the associated trait name and, when needed, the curated lipid class table. Because class-ratio traits can span multiple classes, per-class overlap was stratified by trait layer. Classes were tested only when both trials had at least three candidates.

### Gene Ontology Enrichment

GO biological-process and molecular-function terms came from the PANTHER annotation but were analysed separately, each against sorghum genes. Enrichment was computed within each lipid class ×trait layer ×ontology ×trial, attributing terms to a class’s traits rather than to the full candidate set. Strata with ≥2 candidate genes were tested using a right-tailed hypergeometric test and BH-corrected across all terms in that ontology.

Because LD-mapped candidates are not independent, terms were also tested with a permutation null that preserves the LD block structure. The genome was binned into consecutive 250 kb intervals, and each candidate was assigned to the interval containing its start, so counts reflect distinct regions. Each permutation replaced the observed set of intervals with the same number of gene-density-matched intervals contributing the same number of genes. We used 2,000 draws per term, increasing to 100,000 for terms with fewer than 20 hits, setting the minimum attainable p-value to 1 10^−5^. These p-values were BH-corrected and reported as *q*_*LD*_.

Because GO nesting and shared candidate sets cause repeated signals, terms with identical candidate gene sets were merged, labeled by the most specific term while retaining all merged terms, GO IDs, and lipid classes where the set appeared. This reduced 344 LD-aware significant terms to 217 distinct gene sets across 159 terms. We interpret only gene sets whose term draws on ≥30 background genes and that span ≥2 independent intervals. Of the 87 within the background limit, 71 met the interval requirement. All 217 gene sets are listed in Supplementary Table S2, with the uncollapsed per-term results in Supplementary Tables S3 and S4 and the genes making up each enriched term in Supplementary Table S5, so nothing excluded here is lost.

### Genomic heritability

Genomic heritability was estimated as *h*^2^ = *V*_*g*_ /(*V*_*g*_ + *V*_*e*_) from *y* = **1***µ* + *u* + *e* with 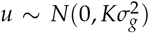, where *K* is a genomic re-lationship matrix built in PLINK 2 (`–make-rel square`) from the LD-pruned GWAS genotypes (`–indep-pairwise 50 5 0.2`) covering 401 accessions.. Variance components were estimated by REML using the eigendecomposition of *K*, reparameterized as *V* = *ε*^2^(*h*^2^ *K* + (1 *h*^2^)*I*) to profile *h*^2^ directly. Estimates use the accessions of each trial present in it: 351 under CTL, 357 under LIN, and the 351 common to both for the paired comparison. Estimates matched `lme4` (Bates *et al*. 2015) within 0.0005 across 34 class-sum traits. We report 95% profile-likelihood intervals. The *h*^2^ was called non-zero when the interval excluded 0.

For per-species comparisons, CTL and LIN were fitted to the same 351 genotypes and 164 shared species, differing only by phenotype. Phenotypic variation per species was summarized as the coefficient of variation of %TIC across genotypes and compared between trials using a paired Wilcoxon signed-rank test. Class sums were fitted for all accessions and species within each trial using summed, spatially corrected intensities transformed as log_10_(*x* + 1). CTL–LIN comparisons reflect trial contrasts, not nutrient-treatment effects.

### Lipid Metabolic Network Analysis (LINEX2)

We used the Lipid Network Explorer (LINEX2; https://exbio.wzw.tum.de/linex/) to reconstruct annotation-guided lipid reaction networks and identify plausible biochemical connections among lipid classes and candidate enzymes. Lipid names were standardized to align with species notation (class plus acyl composition). Since *Sorghum bicolor* is not implemented as a default organism in LINEX2, we selected *Oryza sativa* (OSA) as the reference species for network construction.

For branch-level analysis, we extracted candidate reactions linking the DG/MG, TG, and lysophospholipid pools and summarized each with a reaction-balance score. Three were from the LINEX2 network, while a fourth, an LCAT-like acyl-editing branch denoted LCAT*, was manually defined based on reaction specificity (RHEA:32843, RHEA:44236) rather than retrieved from LINEX2, and the asterisk marks it throughout. For sample *i* and class *c*, define *ω*_*i,c*_ = log_10_(*C*_*i,c*_ + *ε*_*i*_) as the log class-sum abundance, where *ε*_*i*_ is a sample-specific pseudocount.

Each branch score is

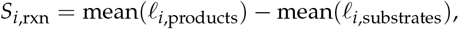

Thus, larger values indicate a product-weighted balance, while smaller values indicate a substrate-weighted one. The substrate and product classes of each branch are given in the supplementary reaction table. These scores are branch-aligned summaries of lipid-pool balance, not measures of enzymatic activity or flux. They were compared between trials by genotype-matched paired Wilcoxon signed-rank tests with BH adjustment.

### Lipid Ontology Enrichment and Hierarchical Classification (LION/web)

We conducted a functional ontology for lipids using LION/web (Molenaar *et al*. 2019) (http://www.lipidontology.com). Lipid nomenclature was standardized according to LIPID MAPS annotations. The default settings of LION/web were applied for enrichment statistics and multiple-testing correction. We retained terms for *q ≥*0.05. A hierarchical classification analysis of individual lipids and functional ontologies was also performed using LION/web.

## Data Availability

All code, figures, and the analysis pipeline are in the GitHub repository at https://github.com/rr-lab/SAP-Lipidomics-Database. Processed lipidomics data, GWAS results, and candidate-gene tables are browsable through the SoLD Shiny application at https://nirwan.shinyapps.io/SAP-Lipidomics-Database/. Raw LC– MS data are deposited in the MassIVE repository under accession MSV000101940 (https://massive.ucsd.edu/ProteoSAFe/dataset.jsp?accession=MSV000101940). SAP genotype data were obtained from Boatwright *et al*. (2022).

## Results

### QC and normalization yielded comparable LC-MS performance across conditions

We profiled early-stage lipids from SAP grown in two field conditions defined earlier. The workflow (Supplementary S1 Fig) comprised tissue collection and extraction, high-resolution LC-MS lipidomic profiling, GNPS-assisted lipid annotation, post-acquisition data correction with SERRF and SpATS (Fan *et al*. 2019; Rodriguez-Alvarez *et al*. 2018), and data dissemination via the SoLD Shiny app (https://nirwan.shinyapps.io/SAP-Lipidomics-Database/).

Run-order quality control showed stable analytical performance (Supplementary S2 FigA–B), with the total ion current (TIC) and the summed signal intensity per sample meeting expected values across system checks, internal standards, and blanks. TIC variability among QC samples was low (7.5% CV for CTL and 6.0% CV for LIN). SERRF normalization improved technical precision, reducing median QC-RSD from 3.6% to 2.0% in CTL and from 1.6% to 0.3% in LIN. The proportion of features with QC-RSD below 30% rose from 75.7% to 94.1% and from 77.3% to 89.3% respectively (Supplementary S2 FigC–D).

PCA of normalized data showed tighter sample clustering than unprocessed data (Supplementary S2 FigE–F). After spatial correction, the leftover variation in MGDG(18:3/18:3) is scattered randomly across the field rather than forming rows, columns, or patches, indicating that position in the field no longer drives the signal (Supplementary S2 FigG–H). Thus, the post-correction lipid signal distributions remained similar across cohorts, and were used for all subsequent analyses.

### Genotypic lipid variation is largely independent of botanical race, and heritable variation is confined to LIN

SAP is structured by botanical race and geographic origin (Casa et al. 2008; Boatwright et al. 2022). Next, we tested whether lipid variation is affected by race or marker-based genetic clusters (*K* = 6; group sizes in Supplementary Table S6). Using Kruskal– Wallis with BH correction within trials (Supplementary Table S7), we found little association. No lipid-class sum was significant for race in either trial. The genetic cluster was significant only for PS under LIN (*q* = 1.16 × 10^−6^, *ε*^2^ = 0.104; Fig. 1). PS is carried by a single species under LIN, PS(18:0/18:2), so the class sum and that species are the same measurement. The same class-level test under CTL, where PS is carried by two species, was not significant (*ε*^2^ = 0.020, *q* = 0.358). Furthermore, none of the species-level tests were significant under CTL, and nine were significant under LIN, eight of them by genetic cluster alone and one, 11,14,17-eicosatrienoic acid, by both genetic cluster and botanical race (Table 1; Supplementary Table S8). PCA of class composition likewise showed minimal structure, with groupings explaining ≤ 1.6% of variation (Supplementary S3 Fig; Sorghum Lipid Database (SoLD): a database curation for lipid information for Sorghum Association Panel Supplementary Table S9). Thus, ancestry explains little of the lipid variation in either trial.

**Table 1.** The individual lipid species associated with ancestry, all of them under LIN. Every species detected in a trial was tested against botanical race and against marker-based genetic cluster (Kruskal–Wallis, Benjamini–Hochberg corrected within trial and grouping). No species reached *q <* 0.05 under CTL for either grouping; the nine below are the complete LIN set. *ε*^2^, effect size; *h*^2^ cond., heritability with the three leading eigenvectors of the relationship matrix fitted as fixed effects; cond. *>* 0, whether the 95% interval still excludes zero. ^‡^Most of this feature’s signal sits in a few accessions near the detection limit, so it is reported for completeness and not interpreted.

| Species | Class | Genetic cluster |  | Botanical race |  | Heritability |  |  |
| --- | --- | --- | --- | --- | --- | --- | --- | --- |
| | | $\varepsilon^2$ | $q$ | $\varepsilon^2$ | $q$ | $h^2$ | $h^2$ cond. | cond. $> 0$ |
| β-Sitosterol | Sterol Lipids | 0.095 | $7.7 \times 10^{-5}$ | 0.011 | 0.625 | 0.676 | 0.644 | yes |
| PS(18:0/18:2) | PS | 0.088 | $1.2 \times 10^{-4}$ | 0.018 | 0.625 | 0.382 | 0.294 | yes |
| 11,14,17-Eicosatrienoic acid,(Z,Z,Z)-<sup>‡</sup> | Fatty Acyls | 0.074 | $6.5 \times 10^{-4}$ | **0.113** | **0.007** | 0.816 | 0.782 | yes |
| Fucosterol | Sterol Lipids | 0.073 | $6.5 \times 10^{-4}$ | 0.013 | 0.625 | 0.404 | 0.322 | yes |
| Cycloartenol acetate | Prenol Lipids | 0.060 | $4.0 \times 10^{-3}$ | 0.031 | 0.597 | 0.688 | 0.662 | yes |
| PC(18:0/20:2) | PC | 0.048 | 0.021 | 0.000 | 0.643 | 0.102 | 0.028 | no |
| PE(18:0/18:3) | PE | 0.047 | 0.021 | 0.000 | 0.671 | 0.358 | 0.350 | yes |
| AEG(o-15:2/15:0) | AEG | 0.046 | 0.023 | 0.043 | 0.433 | 0.392 | 0.290 | no |
| SQDG(16:0/16:0) | SQDG | 0.041 | 0.043 | 0.049 | 0.428 | 0.486 | 0.454 | yes |

**Figure 1.**
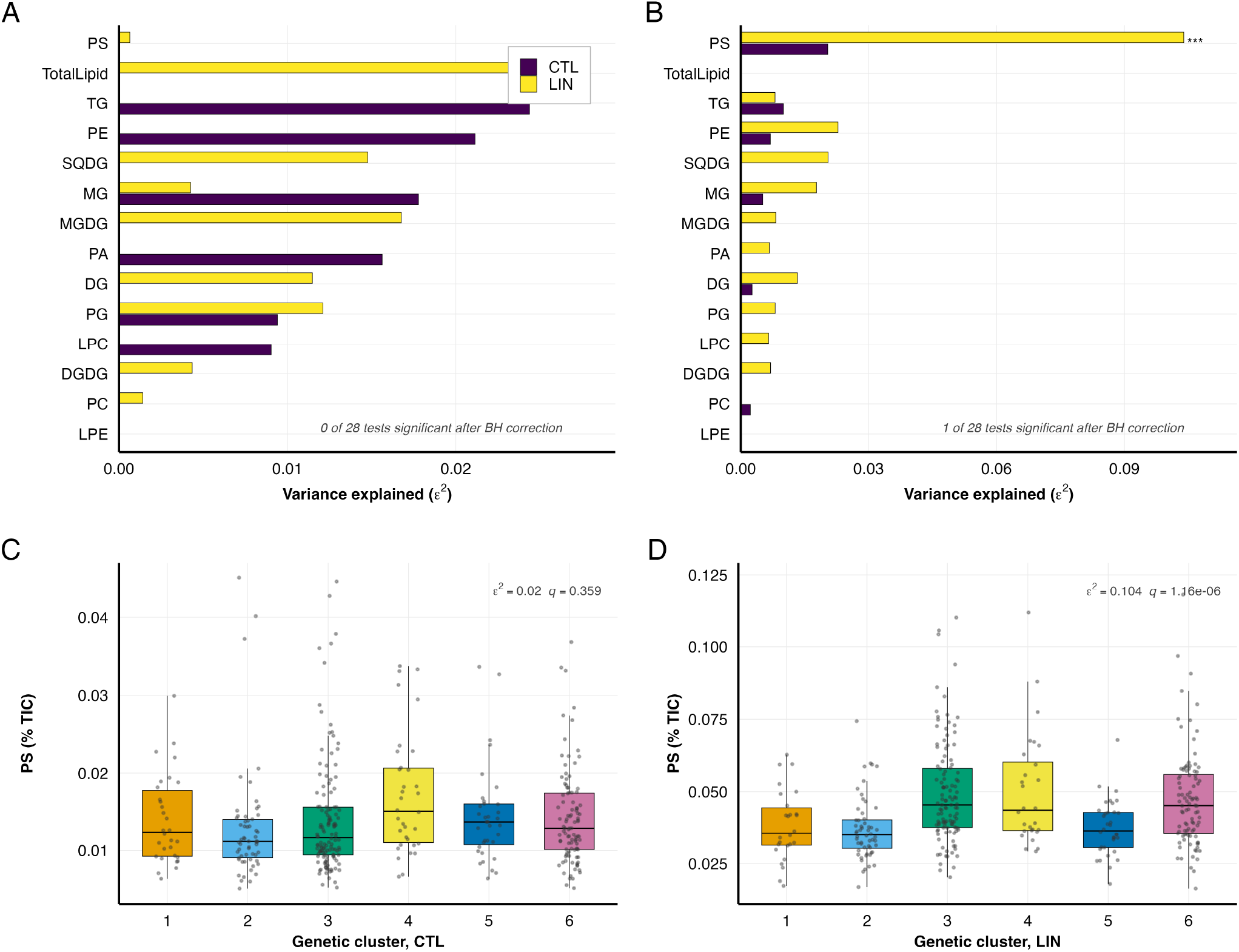
Lipid-class variation is largely independent of botanical race and of population structure, except for PS. **(A**,**B)** Variance explained (*ε*^2^) by botanical race (six groups) and by marker-based genetic cluster (*K* = 6), for the total lipid signal and the 13 lipid classes, computed within each trial. Stars mark tests significant after Benjamini–Hochberg correction. **(C**,**D)** PS abundance (% TIC) by genetic cluster in each trial, each panel on its own scale.

Subsequently, we quantified genetic variance in the lipidome by estimating genomic heritability (*h*^2^) for every species (214 under CTL, 216 under LIN) and compared the trials on the 164 common species measured across the same 351 genotypes (Supplementary Table S10). Under CTL, the median *h*^2^ was near zero, and only 11 of those species had heritability that we could distinguish from zero. Under LIN, the median was 0.171, and 54 species did. The same direction held for species by species comparison, with *h*^2^ higher under LIN for 132 species, higher under CTL for 21, and equal for 11 (Supplementary S4 Fig), and for class by class comparison where every class was higher under LIN except TG and PA (Supplementary Table S11). Estimating *h*^2^ directly on the 17 class sums gives similar results, with a median of 0 under CTL compared to 0.117 under LIN. 15 of the 17 classes are higher under LIN (paired Wilcoxon *p* = 0.007), with the same two exceptions. Only four class sums have an interval excluding zero in either trial (Table 2; Supplementary Table S12). Thus, aggregation does not recover heritability, and the trial difference represents a shift in the whole distribution rather than a handful of strongly heritable classes. The contrast is not an artifact of ancestry or of which genotypes were sampled. It survives conditioning on the three leading eigenvectors of the relationship matrix, dropping each botanical race in turn, and twenty random 80% subsamples of the panel, with LIN exceeding CTL in every run (Supplementary Tables S11 and S13). Nor does it follow from noisier measurement under CTL. Across the same 164 species, total phenotypic variation was comparable between trials (median coefficient of variation 0.28 under CTL against 0.30 under LIN, paired Wilcoxon *p* = 0.29). Thus, the two trials are matched on the accessions actually contributing to *h*^2^.

**Table 2.** Genomic heritability of lipid-class sums, for the classes distinguishable from zero. Estimated by profile likelihood against the marker-based relationship matrix, on the summed spatially corrected intensity of the class (log_10_ scale), using the accessions of each trial present in the matrix. Values are *h*^2^ with the 95% profile-likelihood interval; ^∗^ marks an in-terval excluding zero, the criterion used throughout. Four of the 17 classes meet it in one trial or the other; all 17 are in Supplementary Table S12. *N*, species summed into the class in that trial.

| Class | CTL |  | LIN |  |
| --- | --- | --- | --- | --- |
| | $N$ | $h^2$ | $N$ | $h^2$ |
| FA | 1 | 0.111 [0.000, 0.401] | 1 | 0.404 [0.153, 0.708]* |
| PS | 2 | 0.017 [0.000, 0.243] | 1 | 0.382 [0.167, 0.657]* |
| TG | 67 | 0.217 [0.000, 0.564] | 66 | 0.208 [0.017, 0.525]* |
| PA | 1 | 0.555 [0.252, 0.893]* | 2 | 0.000 [0.000, 0.123] |

Moreover, heritability increases unevenly across classes (Table 3). Under CTL, it is largely confined to TGs, which account for 19 of the 67 TG species and 19 of the 20 heritable species in that trial. Under LIN, it concentrates in the plastid membrane lipids, with 4/5 MGDG, 6/10 SQDG and 4/7 DGDG, the highest proportions of any class with more than one species. MGDG and SQDG also carry the largest absolute shifts in %TIC of any class apart from TG, falling by 2.6 and 1.5 percentage points while TG rises by 2.3, consistent with both the compositional and the genetic changes centering on the photosynthetic membrane lipidome. TG carries heritable species in both trials, though not of the same kind. Every one of the 19 heritable CTL TG species contains an acyl chain of 14 carbons or fewer and sits below 0.05% of TIC, and 14 of the 20 heritable CTL species have a coefficient of variation above one, against 2 of the 66 heritable species under LIN. The CTL set therefore behaves closer to detection or non-detection than to graded abundance, while the 25 heritable LIN TG species are long-chain and vary within the range of the rest of the lipidome. PS, PA, and LPE each include a single species, so they reflect that species rather than the class. The near-absence of heritable species under CTL is consistent with either less genetic variance or a larger residual share, and because neither trial was replicated, we have no independent estimate of plot error to separate the two. The genetic share of variance in the leaf lipidome is therefore much larger under LIN than under CTL, and the GWAS results below follow that asymmetry.

**Table 3.** Heritable lipid species are concentrated under LIN and fall mainly on the plastid membrane classes. Species, the number of that class detected in the trial; med. *h*^2^, the class median; *n>*0, the number whose 95% profile-likelihood interval excludes zero; cond., the number still excluding zero under LIN with the three leading eigenvectors of the relationship matrix fitted as fixed effects. The two counts are not nested, because removing structure can sharpen a borderline signal as well as remove one. Each trial is described as measured; the paired comparison in the text uses only the 164 species common to both. ^†^A single species, so the entry describes that species, not a class. ^§^Fatty acids, ether lipids, sphingolipids and other minor categories.

| Class | Species |  | CTL |  | LIN |  |  |
| --- | --- | --- | --- | --- | --- | --- | --- |
| | CTL | LIN | med. $h^2$ | $n>0$ | med. $h^2$ | $n>0$ | cond. |
| TG | 67 | 66 | 0.084 | 19 | 0.190 | 25 | 28 |
| PC | 36 | 38 | 0.000 | 0 | 0.095 | 7 | 7 |
| Other<sup>§</sup> | 39 | 45 | 0.000 | 0 | 0.128 | 13 | 13 |
| PE | 14 | 15 | 0.000 | 0 | 0.174 | 4 | 2 |
| DG | 16 | 15 | 0.000 | 0 | 0.066 | 0 | 2 |
| SQDG | 11 | 10 | 0.000 | 0 | 0.341 | 6 | 5 |
| DGDG | 7 | 7 | 0.000 | 0 | 0.202 | 4 | 2 |
| MGDG | 5 | 5 | 0.000 | 0 | 0.456 | 4 | 4 |
| MG | 8 | 5 | 0.000 | 0 | 0.098 | 0 | 0 |
| LPC | 4 | 3 | 0.000 | 0 | 0.164 | 1 | 1 |
| PG | 3 | 3 | 0.000 | 0 | 0.158 | 1 | 0 |
| PA<sup>†</sup> | 1 | 2 | 0.554 | 1 | 0.000 | 0 | 0 |
| PS<sup>†</sup> | 2 | 1 | 0.000 | 0 | 0.382 | 1 | 1 |
| LPE<sup>†</sup> | 1 | 1 | 0.000 | 0 | 0.020 | 0 | 0 |
| **Total** | **214** | **216** |  | **20** |  | **66** | **65** |

#### Lipidome overview for CTL and LIN SAP

We quantified a total of 266 lipid species, with 164 detected in both trials, 50 only in CTL, and 52 only in LIN (Supplementary S5 Fig). All lipid values for both trials are browsable in the SoLD Shiny application. TGs and PCs were the most diverse classes in both environments (TG: 67 in CTL, 66 in LIN; PC: 36 in CTL, 38 in LIN; Supplementary S5 Fig; Supplementary Table S14). The largest contributors to between-genotype variance are the most abundant species in both trials, led by MGDG(18:3/18:3) and PC(16:0/18:2) (Supplementary Table S15).

Furthermore, we summarized lipid abundance as %TIC, so the values represent the relative signal in each condition. In both trials, the glycerolipids and glycerophospholipids dominated the annotated signal (CTL: 65.4% and 30.3%; LIN: 64.3% and 30.7%), while the prenol lipids were the only other category above 1% (3.4% and 4.2%) (Fig. 2A). The sterol lipids show the largest proportional change of any category, rising 5.1-fold from 0.04% to 0.22% of the annotated signal, almost all of it β-sitosterol.

**Figure 2.**
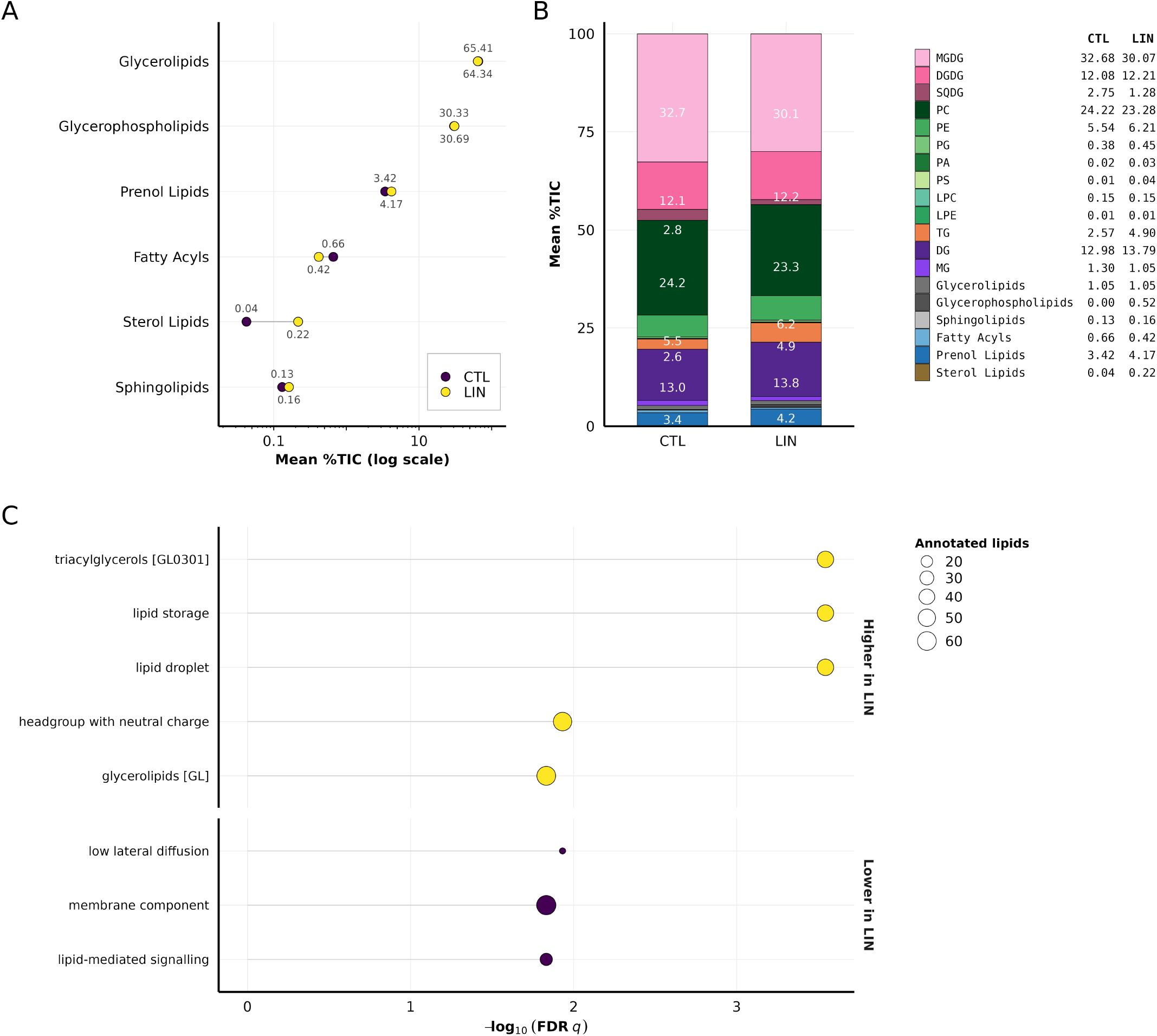
Lipid-class composition of the SAP leaf lipidome in each field trial. Composition is %TIC, the share of total annotated signal, taken per sample and then averaged. **(A)** Annotated LIPID MAPS categories, log scale. **(B)** Annotated species, stacked under their lipid class where they carry a shorthand name and under their LIPID MAPS category otherwise. **(C)** LION terms significant at FDR *q <* 0.05, sized by the number of annotated lipids; the two structural bins among the ten, *PC(38:2)* and *C20:1*, are in Supplementary Table S20 only. Panel C is a direct CTL–LIN contrast and the trials differ in year, field and batch.

Within the galactolipids and phospholipids, MGDG contributed the most with 32.7% (CTL) and 30.1% (LIN), while PC contributed 24.2% and 23.3%. DGDG (12.1%, 12.2%) and DG (13.0%, 13.8%) followed. TG increased from 2.6% (CTL) to 4.9% (LIN), while SQDG decreased from 2.8% to 1.3%. Overall, these thirteen classes comprised 94.7% (CTL) and 93.5% (LIN) of annotated signal. The remainder is carried by species with no shorthand class name, chiefly prenol lipids (3.4% and 4.2%) and fatty acyls, which Fig. 2B shows under their LIPID MAPS category (Supplementary Table S16). Class-level contrasts between trials on the centered and additive log-ratio scales are given in Supplementary Table S17. These class shares do not depend on panel composition, the largest class moving by at most 0.21 percentage points under race-dropping and repeated 20% subsampling (Supplementary Table S18).

Additionally, two minor pools show large changes masked in the bulk profile. PS increases from 0.014% to 0.045% of annotated signal, and its correlation with PC flips between trials (*r* = 0.45 in CTL vs. *r* = −0.34 in LIN; Supplementary Table S19; Sorghum Lipid Database (SoLD): a database curation for lipid information for Sorghum Association Panel Supplementary S6 Fig). Among lysophospholipids, the two classes move in opposite directions. LPE rises from 0.006% to 0.014% of annotated signal while LPC falls on both log-ratio scales (CLR −0.14, ALR −0.62; Supplementary Table S17), so the LPC/LPE ratio halves from 23.2 to 11.1. Both pools stay below 0.2% in both trials.

Within each trial, PCA of the major glycerolipid and glycerophospholipid pools showed lipid class to be the main driver of species-level variation (Supplementary S7 Fig). TG species formed a distinct cluster separated from most membrane lipids in both trials, though along different principal axes, while galactolipids and glycerophospholipids partly overlapped but occupied class-enriched regions, supporting class-level aggregation for the compositional and reaction-network analyses that follow. We next asked whether the class shifts were accompanied by shared structural and biophysical properties using LION, a lipid ontology that groups species into such categories (Molenaar *et al*. 2019). LION ranking analysis across the shared lipid species returned ten significant terms, five higher and five lower with respect to LIN (*q <* 0.05; Fig. 2C; Supplementary Table S20).

Higher in LIN were *triacylglycerols, lipid storage* and *lipid droplet*, all three annotated on the same 44 species, together with *head-group with neutral charge* and *glycerolipids*. Lower in LIN were *the membrane component, low lateral diffusion, lipid-mediated signaling*, and *PC(38:2)* and *C20:1*, the last two relying on too few annotated lipids to interpret (Supplementary Table S20). The five higher in LIN all report the same thing, that storage lipid increased, rising from 16.9% to 19.7% of annotated signal. The three lower in LIN are from the membrane side. *Membrane component* is the bilayer itself, and it falls with the shared phospholipids, from 30.2% to 28.5% of annotated signal, almost all of it PC (24.2% to 21.9%). *Low lateral diffusion* is the tightly packed, ordered fraction of that bilayer, and *lipid-mediated signaling* is the DG, PA, and lyso pool released by membrane turnover, and both are lower in LIN as well. Together, these terms suggest a leaf holding less bilayer and more storage lipid under LIN, consistent with acyl chains being routed into droplets rather than into new membrane. Of these shifts, LION recovers only TG enrichment, as galactolipids and the sulfolipid are missing from the ontology. PS and lysophospholipids have too few shared species for a rank test.

Finally, we summarized each lipid class in a reduced chemical space using the abundance-weighted mean total carbon number and double-bond count across molecular species (Supplemen-tary S8 Fig; Supplementary Table S21). Carbon number cleanly separates the classes, from lysophospholipids and monoacyl-glycerols through diacyl lipids to TG, and diacyl unsaturation varies independently of chain length. Across trials, PE shows the largest unsaturation increase among classes with more than ten species; otherwise, aside from a higher TG carbon number, every class except the non-comparable PS changes by less than one carbon and 0.25 double bonds. Within-class chemistry is therefore largely stable, and CTL–LIN differences reflect shifts in class proportions rather than chain composition.

Compared with CTL, four population-wide shifts characterize the LIN lipidome: (i) SQDG depletion, (ii) TG enrichment, (iii) a phospholipid rebalancing centered on PS, and (iv) an LPC/LPE balance shifted toward LPE. All rests on the abundance and composition analyses. LION corroborates only the TG shift. Because the trials also differ by year, field, planting date, and analytical batch, these shifts characterize LIN as a whole and cannot be attributed to just nutrient supply and cold. The reaction-network and GWAS sections that follow ask which reactions and which loci accompany them. Candidate genes below follow for the TG and SQDG shifts. The PS and lysophospholipid changes sit on pools under 0.2% of TIC and are not resolved to candidates, although PS is the one class whose abundance tracks the genetic cluster (Fig. 1C,D).

#### GWAS recovers more genomic regions under LIN and more trait-sharing per region under CTL

We ran separate GWAS for each individual lipid trait, lipid class sum, and lipid class ratio in both trials (747 in total: 220 individual species and 153 class sums and ratios under CTL, 221 and 153 under LIN), using a univariate linear mixed model with the first three genotype PCs to account for population structure (Fig. 3). SNPs surpassing the Bonferroni genome-wide significance threshold (*p* ≤ 8.19 10^−9^ for CTL and 8.22 × 10^−9^ for

**Figure 3.**
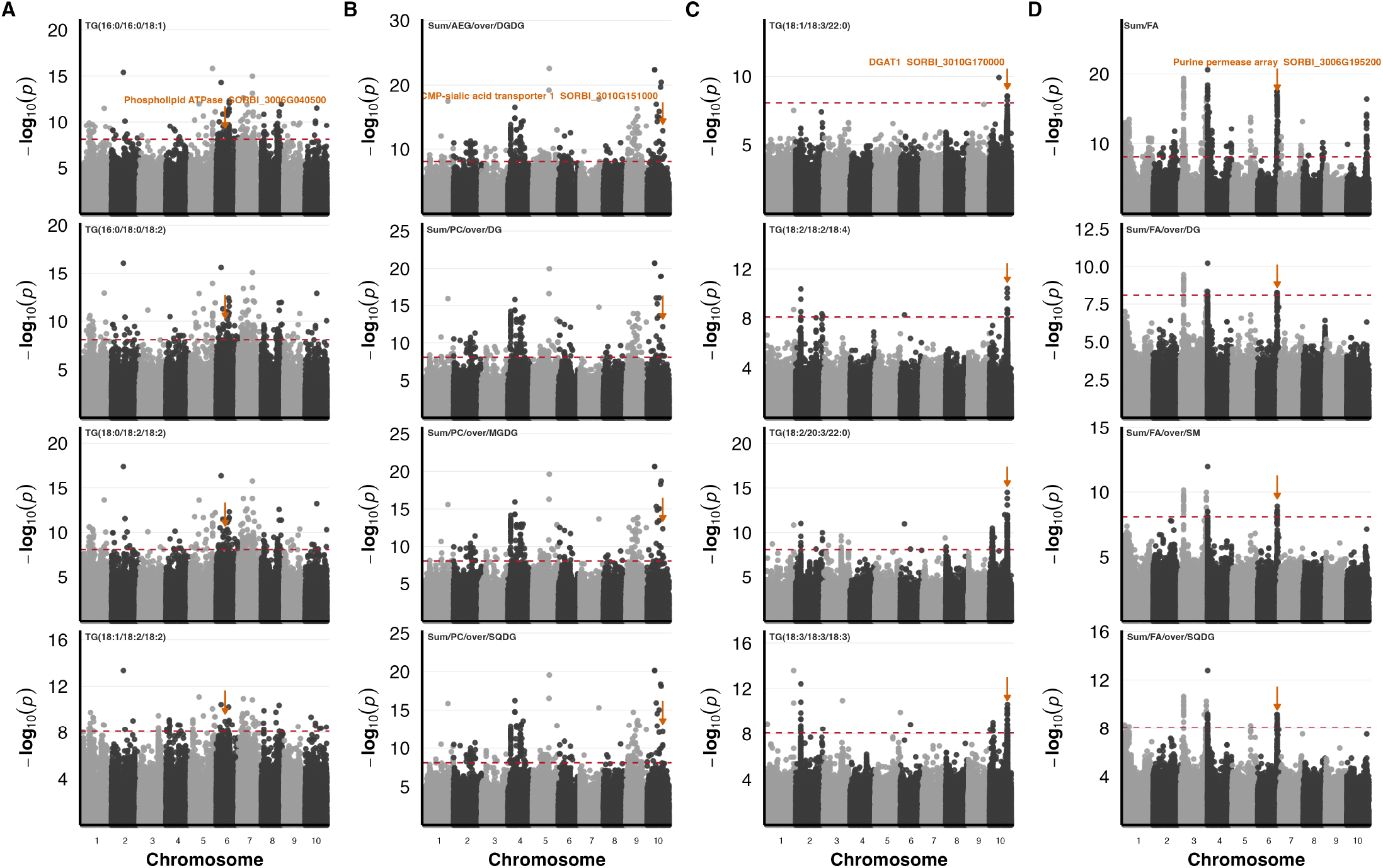
The same candidate locus recurs across multiple lipid phenotypes within each trial. Each column stacks four phenotypes sharing one candidate locus. An orange arrow marks the lead SNP, the gene is named once above it in the top plot, and each plot is labelled by its trait. **(A)** CTL individual lipids, `SORBI_3006G040500` (phospholipid-transporting ATPase, chr 6). **(B)** CTL class sums and ratios, `SORBI_3010G151000` (CMP-sialic acid transporter 1, chr 10). **(C)** LIN individual lipids, `SORBI_3010G170000` (*DGAT1*, chr 10). **(D)** LIN class sums and ratios, the chr 6 purine-permease array (`SORBI_3006G195200`). Red dashed lines, Bonferroni genome-wide threshold (*p* ≥ 8.19 *×* 10^−9^); point shading alternates by chromosome.

LIN) were mapped to candidate genes using the LD-based cutoff of 0.4. Under CTL, this identified 1,062 candidate genes for individual lipids and 54 for class sums and ratios (Supplementary Table S22). Under LIN, the same procedure identified 4,367 and 385 candidate genes (Supplementary Table S23).

Candidate gene counts do not equal signal counts because one significant SNP tags all genes in its LD window. After binning candidates into 250 kb windows, there were 358 distinct windows under CTL and 948 under LIN. So, LIN captures ~ 2.6 × more loci, not the four-fold suggested by gene counts. Hence, CTL loci average 3.0 candidate genes, while LIN loci average 4.6. Candidates also differ in phenotype recurrence. In the individual-lipid layer, 586 of 1,062 CTL candidates were linked to more than one phenotype compared to 1,315 of 4,367 under LIN. Thus, CTL yields fewer loci, each carrying many lipid associations, while LIN yields more loci carrying few. The class-sum and ratio layer is too small under CTL for the same contrast (Supplementary Table S22). Because lipid phenotypes are highly correlated by design, these recurrence counts are descriptive rather than formal tests against a null; the cross-trial comparison in the next section is evaluated explicitly.

Table 4 lists the five loci linked to the most phenotypes in each trial and trait layer, one row per signal with the number of genes it tags. The trial difference is not just a counting artifact, since genomic heritability is also higher under LIN, and that estimate uses no LD windows or gene counts. Traits with higher heritability are easier to map. Therefore, the longer LIN candidate list follows from the heritability difference rather than Sorghum Lipid Database (SoLD): a database curation for lipid information for Sorghum Association Panel confirming it.

**Table 4.**
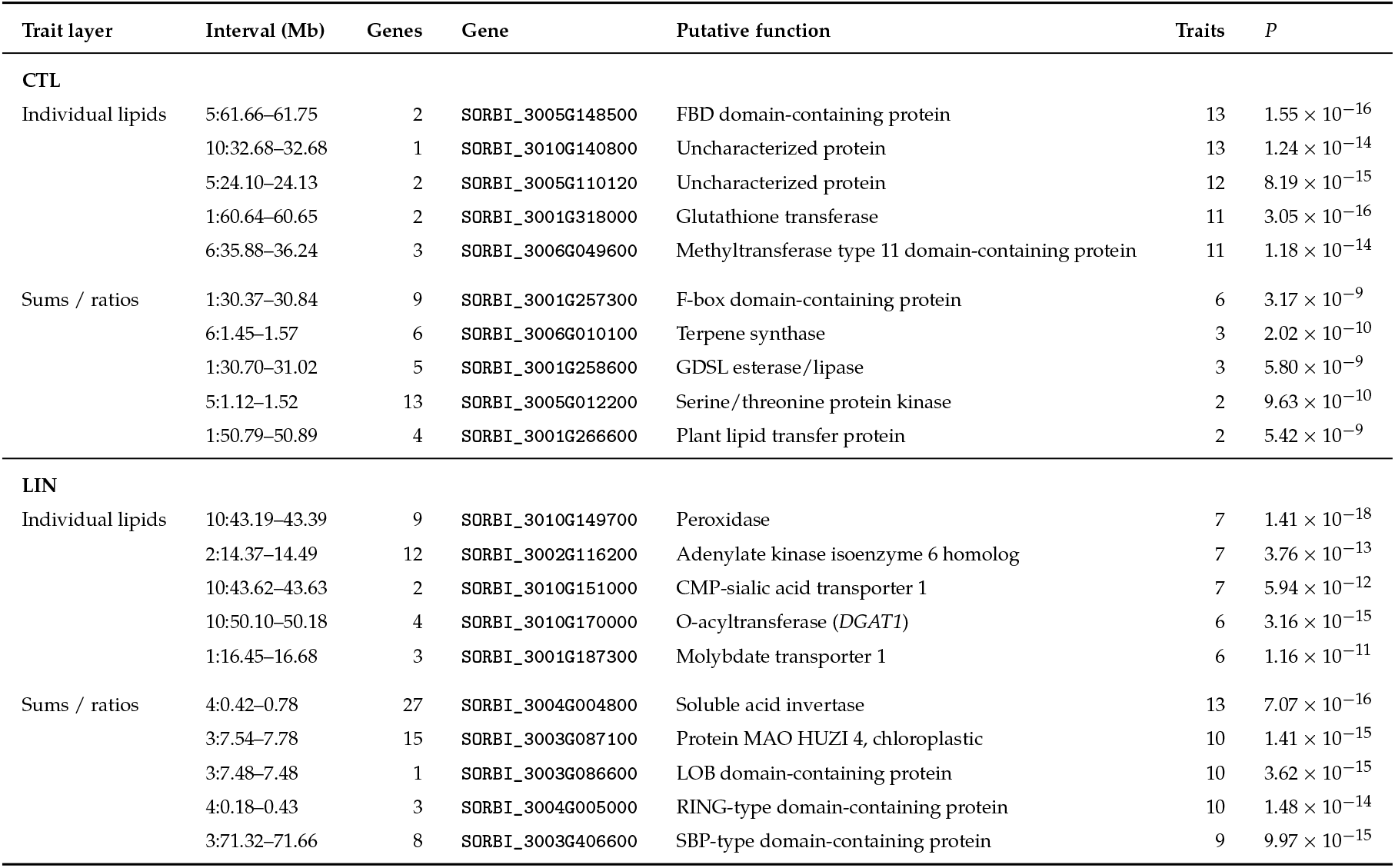
The most recurrent GWAS loci under CTL and LIN. The five loci associated with the largest number of phenotypes in each trial and trait layer. A locus is one signal, defined as all candidate genes sharing a trial, trait layer, chromosome and best *p*-value, so the genes tagged by one lead SNP occupy a single row. Interval, the span of the tagged genes in Mb; Genes, the number of candidate genes the signal tags; Gene, the first annotated gene in that interval, given as a label for the locus rather than as a causal assignment; Traits, the number of lipid phenotypes associated with the locus. Recurrence counts are descriptive summaries of the candidate tables and not a test against a null expectation, since lipid phenotypes are correlated by construction.

#### LD-aware GO enrichment reduces the candidate lists to a small number of interpretable terms

Both candidate lists were too large to inspect gene by gene, so we tested Gene Ontology (GO) biological-process and molecular-function terms for over-representation within each trial lipid class, using a permutation null that preserves LD-block structure. This distinguishes terms supported by multiple independent intervals from those driven by a single LD block.

In total, 344 of 1,292 tested terms passed the LD-aware null (*q*_LD_ *<* 0.05). Because GO terms are nested and class sums and ratios share candidate genes, we merged terms with identical gene sets, leaving 217 term–gene-set combinations across 159 distinct terms (Supplementary Table S2). Of these, 87 draw on ≥ 30 background genes, while the rest were too broad to interpret such as *zinc ion binding* (909 genes) and *transmembrane transport* (865 genes). Seventy-one of the 87 are supported by two or more independent intervals, while the remaining 16 rest on a single LD block. Figure 4 shows the strongest four per lipid class among the 71, giving 22 gene sets.

**Figure 4.**
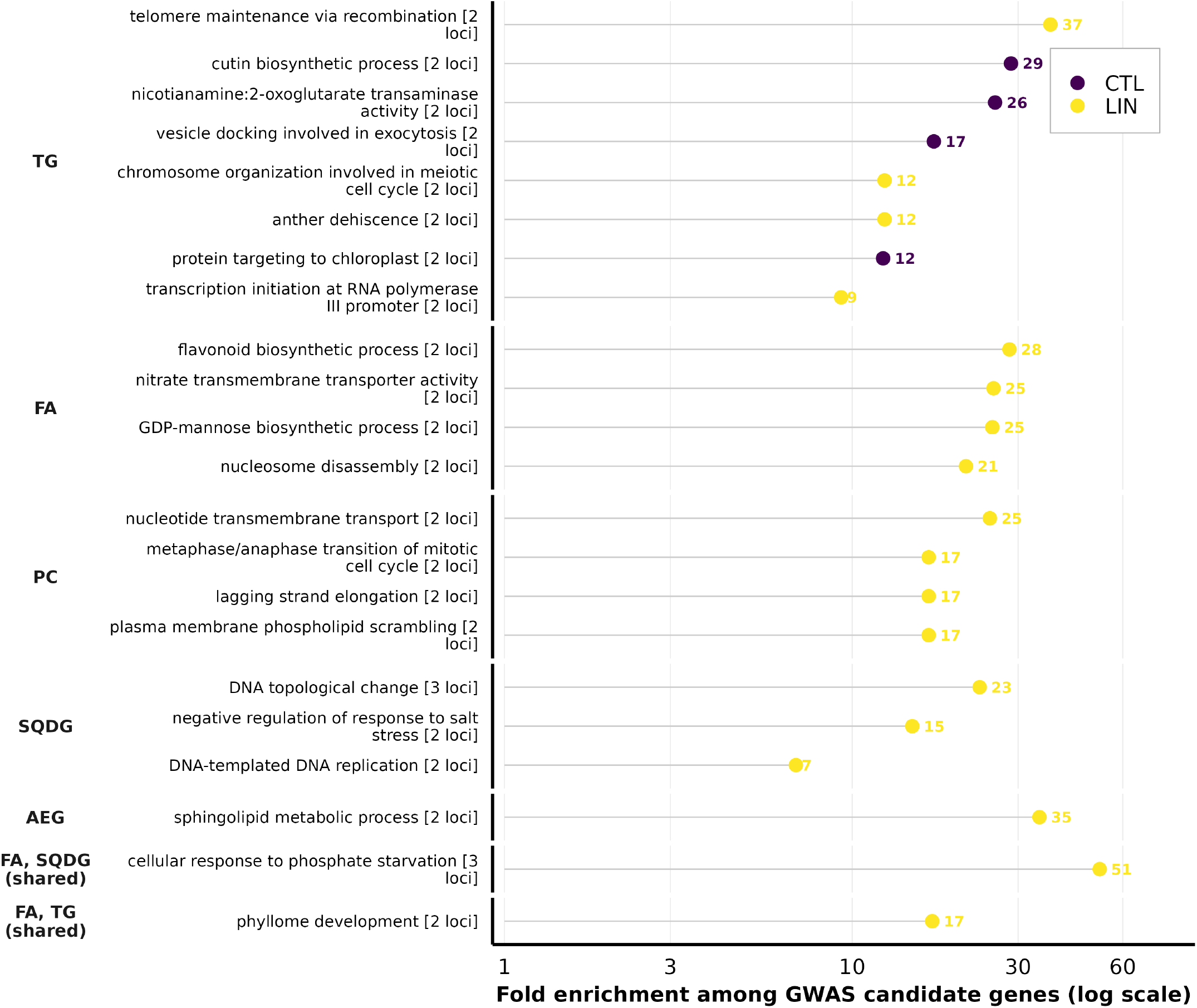
Gene Ontology terms enriched among LD-mapped GWAS candidate genes, grouped by the lipid class whose traits produced them. Twenty-two gene sets are shown, the strongest four per lipid class among the 71 that pass the LD-aware permutation null (*q*_LD_ *<* 0.05), draw on at most 30 background genes and span at least two independent intervals. The background limit is a criterion of interpretability rather than significance, since terms such as *zinc ion binding* (909 background genes) or *transmembrane transport* (865) pass the null without naming a lipid-related process; the interval requirement removes 16 gene sets resting on a single LD block. Points give fold enrichment over chance on a logarithmic axis and are coloured by trial, and each label carries the number of independent intervals supporting it. Left-hand strips name the lipid class; the two marked shared are gene sets recurring across several classes, which are correspondingly less specific.

Those 22 gene sets are unevenly split between trials, with 18 under LIN and 4 under CTL, the same asymmetry that the candidate counts show. All four CTL gene sets come from TG traits. The LIN gene sets center on nutrient transport and signaling. *Cellular response to phosphate starvation* is the strongest of them, at 51 × enrichment across three independent intervals, and is one of only two gene sets with more than one lipid class. *Nitrate transmembrane transporter activity* follows from fatty-acid traits at 25 × across two intervals, and *sphingolipid metabolic process* from AEG at 35 ×. Fatty acids, PC, and TG each contribute four gene sets, and SQDG contributes three. Enrichment therefore narrows the 5,173 candidate genes to the 919 that fall within an enriched term, a short list that could be used for functional validation.

#### Candidate genes overlap between trials only for individual species, and the overlap does not survive collapsing linkage

The two trials recovered largely different candidate lists, but different GO terms cannot indicate whether the underlying genetics differ. We therefore counted how many candidate genes the two trials share and compared that count with the chance expectation from the 34,027 *Sorghum bicolor* v3.1 gene models, both at gene resolution and after collapsing linked genes into 100, 250, and 500 kb windows Supplementary S9 Fig). Overlap was tested with a hypergeometric test and summarized as fold enrichment over chance and as the Jaccard index, the proportion of all candidate genes appearing in both conditions.

At the gene level, the two candidate sets overlapped more than chance, with 291 genes shared against 140 expected (2.08, *p* = 1.1 × 10^−35^), but the excess comes entirely from the individual-lipid layer (2.07 ×, *p* = 2.4 10^−34^; Table 5). The class-sum and ratio layer shares no gene at all between trials, against 0.6 expected by chance, so it is neither enriched nor depleted, only empty. The shared genes are in any case a small fraction of the total, with a Jaccard index of 0.056. Within the individual-lipid layer, the excess is carried by TG alone (1.74 ×, *q* = 5.3 × 10^−3^), while PC and the fatty acids overlap no more than chance, and four further classes share no genes at all, including SQDG, despite its 400 LIN candidates (Table 6; Supplementary Table S24).

**Table 5.** Cross-trial overlap of GWAS candidates at gene and locus resolution. Overlap between CTL and LIN candidate sets, tested against a background of 34,027 annotated gene models (gene level) or of all genome windows containing at least one gene (locus level). Fold, observed/expected shared; *J*, Jaccard index; *P*, hypergeometric (right-tail) test. Bold rows are the resolutions at which LD blocks are effectively collapsed.

| Resolution | Trait layer | CTL | LIN | Shared | Exp. | Fold | $J$ | $P$ |
| --- | --- | --- | --- | --- | --- | --- | --- | --- |
| Gene | All layers | 1091 | 4373 | 291 | 140.2 | 2.08 | 0.056 | $1.1 \times 10^{-35}$ |
| Gene | Individual lipids | 1062 | 4367 | 282 | 136.3 | 2.07 | 0.055 | $2.4 \times 10^{-34}$ |
| Gene | Class sums / ratios | 54 | 385 | 0 | 0.6 | — | 0.000 | 1 (n.s.) |
| 100 kb | Individual lipids | 494 | 1554 | 185 | 158.5 | 1.17 | 0.099 | $4.4 \times 10^{-3}$ |
| 100 kb | Class sums / ratios | 23 | 118 | 0 | 0.6 | — | 0.000 | 1 (n.s.) |
| **250 kb** | **Individual lipids** | 351 | 947 | 153 | 145.9 | **1.05** | 0.134 | **0.22** |
| 250 kb | Class sums / ratios | 17 | 72 | 0 | 0.5 | — | 0.000 | 1 (n.s.) |
| **500 kb** | **Individual lipids** | 275 | 634 | 146 | 140.7 | **1.04** | 0.191 | **0.26** |
| 500 kb | Class sums / ratios | 13 | 50 | 0 | 0.5 | — | 0.000 | 1 (n.s.) |

**Table 6.** Per-class cross-trial overlap of individual-lipid GWAS candidates. Genes were assigned to a lipid class through the phenotypes with which they were associated.

| Class | CTL | LIN | Shared | Fold | *q* |
| --- | --- | --- | --- | --- | --- |
| TG | 832 | 892 | 38 | 1.74 | $5.3 \times 10^{-3}$ |
| Fatty acid | 105 | 1964 | 6 | 0.99 | 1 (n.s.) |
| Terpenoid | 44 | 38 | 0 | — | 1 (n.s.) |
| PC | 39 | 1319 | 4 | 2.65 | 0.22 (n.s.) |
| DG | 11 | 31 | 0 | — | 1 (n.s.) |
| SQDG | 8 | 400 | 0 | — | 1 (n.s.) |
| AEG | 3 | 122 | 0 | — | 1 (n.s.) |

This mainly reflects LD, because each association maps every gene in its window, and gene-dense regions are mapped repeat-edly in both trials. Because no window size is objectively correct for collapsing linkage, we repeated the test at 100, 250, and 500 kb and read the trend across them as the result. For individual lipids, the excess fell at every step, from 2.07 × at gene level to 1.17 × at 100 kb and 1.05 × at 250 kb, where it was no longer significant (*p* = 0.22), and 1.04 × at 500 kb (*p* = 0.26), so loci recovered in the two trials are no more concordant than two independent samples of the same genome. The class-sum and ratio layer share no window at any of the three sizes.

Recovering the same gene never meant recovering it for the same lipid. Of the 291 shared genes (Supplementary Table S25), none are candidates for the same trait in both trials, although 13 traits have significant associations in both. At locus resolution, the two trials therefore recover essentially independent sets, and the gene-level excess reflects linkage rather than shared genetic control.

#### LINEX-based reaction mapping, reaction-balance scoring, and GWAS support

To place lipid associations in a biochemical context, we used LINEX2 (Rose *et al*. 2023), which integrates curated lipid-reaction data with measured lipid abundances to identify biochemical connections among lipids and enzymes (Morgat *et al*. 2016). Since LINEX2 does not provide a sorghum reference, we used rice instead. Therefore, the resulting network is considered a set of testable hypotheses. The resulting network (Fig. 5A) revealed four candidate branches linking DG/MG, TG, and lysophospholipid pools: (i) PC+DG*⟷*LPC+TG (LCAT*), (ii) TG*⟷*DG (PNPLA1-labeled li-pase branch), (iii) PE+DG*⟷* TG+LPE (LRO1/PDAT-like transfer), and (iv) DG+MG TG (PNPLA3-labeled branch). LINEX2 returned an enzyme assignment for three of these but not for branch (i), so the LCAT assignment was made based on the reaction’s specificity (RHEA:32843, RHEA:44236) and is marked LCAT* throughout. The substrates, products and RHEA identifiers of each branch are given in Supplementary Table S26. These annotation-based hypotheses do not establish *in vivo* flux direction, and because CTL and LIN were sampled in different years, all LIN–CTL contrasts are genotype-matched between-trial ratios rather than causal effects of management.

**Figure 5.**
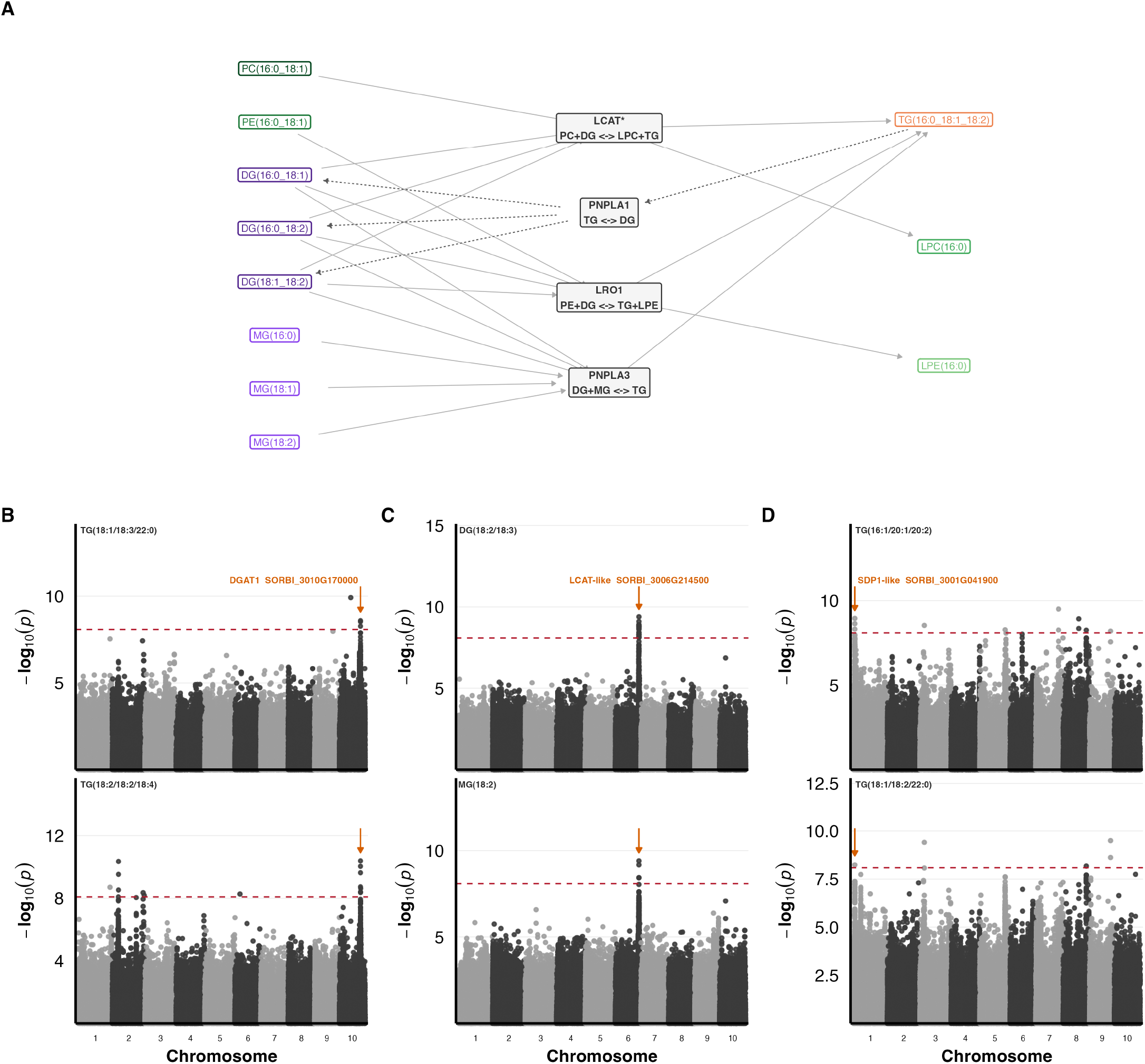
Candidate lipid reaction branches and the GWAS support for them. **(A)** LINEX-enriched subnetwork linking four candidate reactions to their substrate and product species, LCAT-like acyl editing, PNPLA1-mediated TG *↔*DG lipolysis, LRO1-mediated PE/DG*↔* TG/LPE transfer, and PNPLA3-associated DG/MG*↔* TG re-esterification. Arrows show reaction direction, dashed arrows run right to left, and boxed labels give representative species coloured by class. All four reactions came from the LINEX2 network; the label on the acyl-editing branch, marked LCAT*, was assigned from reaction specificity rather than returned by LINEX2. **(B–D)** GWAS support for the three branch-supporting LIN candidates, two traits per gene. **(B)** *DGAT1* (`SORBI_3010G170000`), **(C)** an LCAT-like gene (`SORBI_3006G214500`), **(D)** an SDP1-like lipase (`SORBI_3001G041900`). Plotting conventions as in Fig. 3. The reaction-balance scores are given in Supplementary Table S26 rather than here, because they are a CTL–LIN contrast.

For each branch, we computed a per-sample reaction-balance score as the mean log abundance of products minus substrates (Rose *et al*. 2023; Gieger *et al*. 2008; Illig *et al*. 2010), so a batch effect that scales every lipid by the same factor cancels. In genotype-matched paired Wilcoxon tests across 355 genotype pairs, all four branches shifted, and all four kept their direction when recomputed on only the species measured in both trials. The results were positive for PNPLA3 (+0.314), LRO1 (+0.261), and LCAT* (+0.139), and negative for PNPLA1 (−0.256) (BH-adjusted *p <* 10^−52^; Supplementary Table S26). TG-producing branches shifted in one direction and the hydrolytic TG *⟷*DG branch in the other, showing that the class-level shift holds within genotypes and not just on average. Because substrates and products are different lipid classes, scale invariance does not remove class-specific trial artifacts, so these directions remain trial differences.

Cross-referencing branches with the LIN GWAS candidate tables supported three of four (Supplementary Tables S23 and S27). Branch (i) was supported by two LCAT-superfamily genes, `SORBI_3006G214500` and `SORBI_3004G341900`, whose closest Arabidopsis homologues are LCAT4 and PSAT. The characterized members of this family release a lysophospholipid from PC or PE rather than transferring the acyl group to diacylglycerol (Chen et al. 2012; Banaś et al. 2005). Branch (ii) was sup-Sorghum Lipid Database (SoLD): a database curation for lipid information for Sorghum Association Panel ported by `SORBI_3001G041900`, a PNPLA-domain lipase in the SDP1 clade associated with three TG species under LIN. Finally, branch (iv) by `SORBI_3010G170000` (*DGAT1*). No candidate was found for branch (iii), and no PDAT-family gene was recovered in either trial (Supplementary Table S28). The same cross-reference against the CTL candidate tables returned a sin-gle gene on branch (ii), `SORBI_3005G144300`, a patatin-domain lipase (*p* = 3.10 *×* 10^−10^;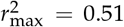). All four of its associated traits are TG species that lie on that branch, against 3 of 12 for the seven LIN genes mapped there, and it also falls in the CTL-enriched term *triacylglycerol lipase activity* (10.7 ×across two independent intervals; Supplementary Table S4), so enrichment and the reaction network recover it independently. Together, these results link the class-level composition shift to four candidate reactions, three of which carry a genome-wide association under LIN and one of those also under CTL.

#### An interactive SoLD Shiny application enables exploration of lipid and GWAS results

The SoLD web server provides an integrated environment for exploring lipid phenotypes and their genetic associations across two trials (Fig. 6A). Traits can be examined as individual species, class-level sums, and inter-class ratios (Fahy *et al*. 2009), with hierarchical filtering by LIPID MAPS category, main class, and species. Exploratory and multivariate modules (data preview, boxplots, volcano plots, correlation and network views, heatmaps, PCA, t-SNE/UMAP) are paired with two genetic modules: GWAS for trait-specific associations and Gene Hits for recurrent candidate genes under user-defined dataset, trait-type, lipid-class, and LD settings. A Genotype module identifies accessions within specified lipid value ranges for experimental design.

**Figure 6.**
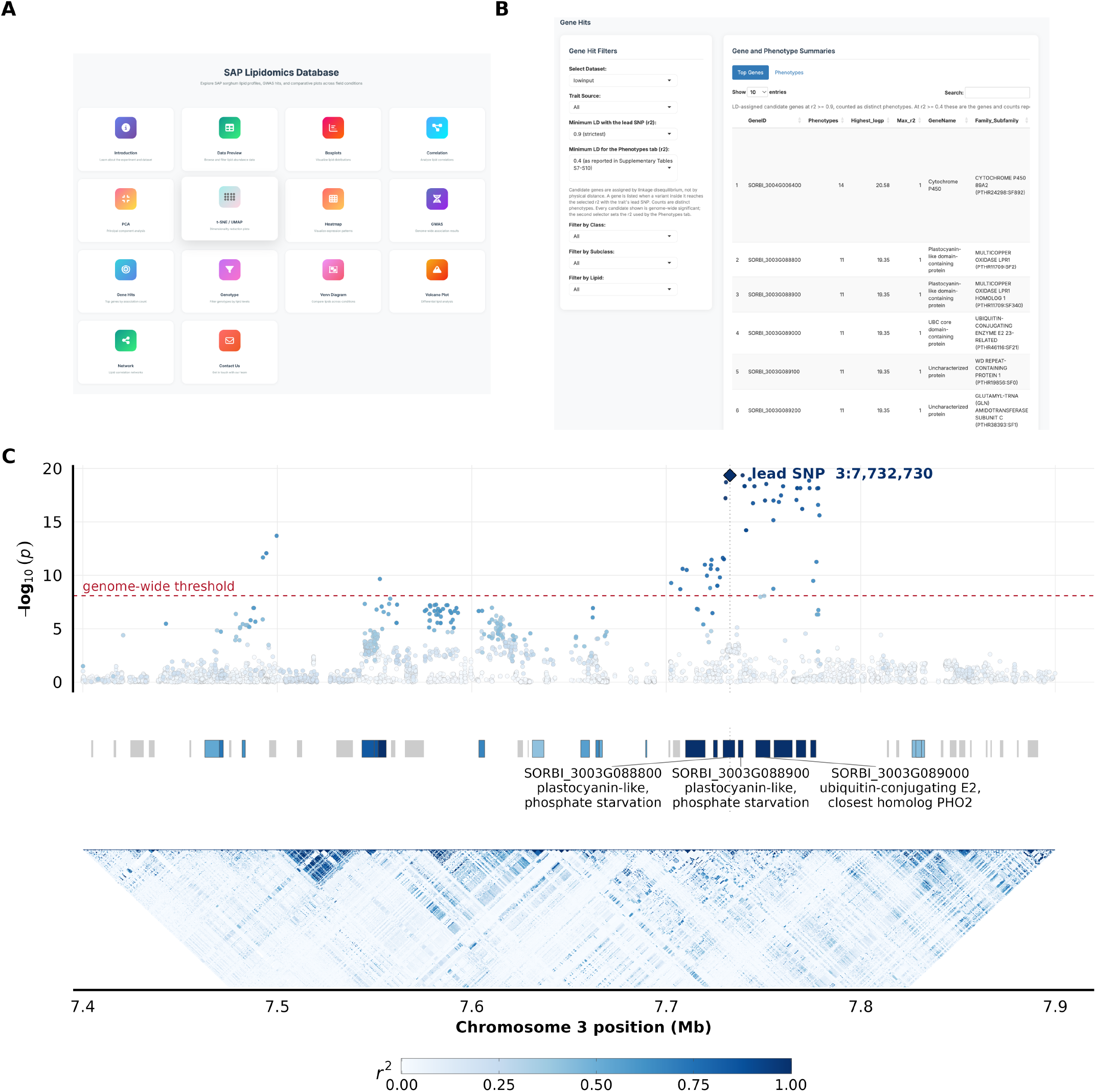
SoLD Shiny app and the chromosome 3 candidate block under LIN. (A) The homepage of the SoLD Shiny app with a list of features. (B) Gene Hits module under the LIN dataset, all trait sources, and the strictest LD setting (r^2^ ≥ 0.9), every gene shown being genome-wide significant, ranking candidate genes by the number of distinct phenotypes each recurs in. (C) Regional view of the chromosome 3 locus that the top of that table points to, over 7.40–7.90 Mb for the LIN fatty-acid class sum. Top, every GWAS SNP as log_10_(*p*), coloured by its r^2^ with the lead SNP, with the LIN Bonferroni threshold dashed and the lead SNP as a diamond; a second SNP at 3:7,739,242 ties it at *p* = 4.46 × 10^−20^. Middle, the 49 annotated genes in the window, coloured by the highest r^2^ any variant inside them reaches with the lead SNP, which is how candidate genes are assigned throughout; grey genes carry no variant at r^2^ ≥ 0.4. Three of the five block genes are labelled, the other two being uncharacterised. Bottom, pairwise r^2^ between variants less than 150 kb apart. All three tiers share one r^2^ scale.

Also, the interface allows for easy tracing of traits to candidate genes through the combination of filtering, plotting, and gene summaries. In the *Gene Hits* module (LIN dataset; all trait Sorghum Lipid Database (SoLD): a database curation for lipid information for Sorghum Association Panel sources; strict LD, *r*^*2*^ ≥ 0.9; every gene shown genome-wide significant), 1,291 candidate genes are ranked by the number of phenotypes in which they recur (Fig. 6B). The top gene is a chromosome 4 cytochrome P450 (`SORBI_3004G006400`), recurring in 14 phenotypes. Next are five adjacent chromosome 3 genes, each carrying a variant in perfect linkage disequilibrium (*r*^2^ = 1) with the most significant SNP at the locus, recurring in the same 11 phenotypes, one class sum (free fatty acids), the species FA(22:1), and nine class ratios, sharing the same association (*p* = 4.46×10^−20^). Within the 500 kb window, the associa-tion concentrates in 7.70–7.78 Mb (Fig. 6C). Three of the five block genes are annotated as two multicopper oxidases of the LOW PHOSPHATE ROOT family and a PHO2-family ubiquitin-*q*, Benjamini–Hochberg-adjusted hypergeometric *P*-value. n.s., not significant. conjugating E2, so the block carries genes of the phosphate-starvation response. LD cannot distinguish among the five, and the associations are with fatty-acid traits rather than phospholipid abundances. Thus, the app lets readers explore the full candidate space by choosing their dataset, trait type, lipid class, and LD threshold; tracing any lipid to its associations and candidate genes not discussed in this paper.

## Discussion

The trials differed in lipid class composition, and their lipid-trait loci were largely distinct. We relate lipidome changes to trial conditions, highlight candidate genes, and conclude with the near-absence of shared loci between trials. GWAS returned more candidates than can be interpreted: 1,062 genes under CTL and 4,370 under LIN for individual lipids alone. Therefore, the genes discussed below are a selection, and the basis for it should be explicit. We applied two filters. A gene set had to survive the LD-aware permutation null on at least two independent 250 kb intervals while drawing on no more than 30 background genes, which leaves 20 sets under CTL and 51 under LIN, and a gene had to fall on one of the four LINEX reaction branches. The background cap is a specificity criterion rather than a significance one, since a term spanning hundreds of genes can only reach shallow enrichment and names no lipid-related process. Within what survived, we then looked for functions related to the ways LIN departed from standard practice, N, P, cold, and the absence of chemical treatment. Recurrence across lipid traits is reported in Table 4, but is not used to select candidates because lipid phenotypes are correlated by construction, and a high trait count reflects that correlation as much as anything about the gene. The two filters ask different questions and return largely different candidate genes to validate. Genes meeting neither are not ranked lower, only unexamined, and the full candidate tables are supplied in Supplementary Tables S19 to S22 and in the Shiny application so that other selections can be made.

### Integrated interpretation of lipidome remodeling across contrasting field trials

The two most prominent changes in LIN are a decrease in SQDG and an increase in TG. The accumulation of TG is commonly associated with low temperatures, drought, oxidative stress, and nutrient-driven carbon reallocation, suggesting a shift towards increased storage within the LIN signature (Tan et al. 2018; Yan *et al*. 2025; Pfaff *et al*. 2020). The median LIN values are higher than CTL for every lipid ratio in which TG is the numerator. Thus, TG rises against every class it was tested against (Supplementary Table S29). DG rises less than TG but more than MG, giving DG/MG 1.3×, while DG/PS falls to 0.32× because PS rises further still. Correlations also change for PS. PS–TG switches from negative in CTL (*r* =−0.26) to positive in LIN (*r* = +0.43), a strong sign reversal (Supplementary Table S19). Additionally, LINEX reactions add to these results. The three TG-producing branches (i), (iii), and (iv) shift towards products under LIN, while TG *↔*DG hydrolysis on branch (ii) shifts towards the substrate, collectively favoring TG over DG (Supplementary Table S26). These class-level balances together suggest a reorganization of DG supply and TG synthesis and mobilization, though direct measures of enzyme activity are needed. Both arms carry LIN candidates. On branch (iv) (DG+MG*⟷* TG), *DGAT1* (`SORBI_3010G170000`) acylates diacylglycerol to triacylglycerol and falls within a repeatedly mapped chromosome 10 crude-fat QTL in sorghum (Boyles *et al*. 2017; Yen *et al*. 2008). On branch (ii) (TG*⟷* DG), the reverse hydrolytic step has the candidate `SORBI_3001G041900`, an SDP1-clade lipase. In Arabidopsis, SDP1 hydrolyzes triacylglycerol to diacylglycerol and free fatty acid and accounts for most TG breakdown during reserve mobilization (Kelly *et al*. 2011). This locus clusters with *SbSDP1-L* in the sorghum lipase phylogeny (Sapara *et al*. 2024). The same reaction network places two LCAT-superfamily acyltransferases (`SORBI_3006G214500`, `SORBI_3004G341900`) on branch (i) (PC+DG *⟷*LPC+TG), which releases lysophospholipids from PC, although characterized members of this family release a lysophospholipid from PC or PE rather than transferring the acyl group to diacylglycerol, so the branch assignment remains a hypothesis (Chen *et al*. 2012; Banaś *et al*. 2005).

Another change involves the balance among phospholipid headgroups. PS increased against every class it was tested against (Supplementary Table S29), and across genotypes, the PS–PC correlation flipped from positive in CTL (*r* = +0.45) to negative in LIN (*r* = −0.34), the largest shift among class pairs. Thus, the two headgroups move together across the panel under CTL but trade off against each other under LIN (Supplementary Table S19). PE showed the largest increase in mean double-bond number of any class, although the species detected differ slightly between trials, and this is a composition measure rather than a direct account of acyl editing. Comparable shifts in headgroup proportions, membrane unsaturation, and lysophospholipid pools are reported under cold, nutrient, oxidative, and water limitation, where they accompany changes in membrane fluidity, lipid repair, signaling, and remobilization in other studies (Hou *et al*. 2016; Rawat *et al*. 2021; Yan *et al*. 2025; Zhang *et al*. 2020). The balance between the lysophospholipid classes also shifted, with LPC/LPE dropping to 0.48-fold in LIN (Supplementary Table S29), driven by rising LPE while LPC remained essentially unchanged (Supplementary Table S16). PS and the lysophospholipids each contribute under 0.2% of TIC, so these are large proportional changes on small pools. The annotation is also shallow, with two PS species under CTL and one under LIN, four LPC species under CTL and three under LIN, and a single LPE species in both, so none of these shifts can be read as evidence of phospholipase activity.

These changes do not indicate phosphate conservation, nor a headgroup swap. Under phosphate limitation, phospholipids are degraded to release phosphate and are replaced by glycolipids, raising SQDG/DGDG and lowering phospholipids (Essig-mann et al. 1998; Härtel et al. 2000; Jouhet et al. 2004; Kelly and Dörmann 2002; Pfaff et al. 2020). However, neither occurs here. On the CLR scale, SQDG drops the most, galactolipids drop relatively less, and PG shows no change (Supplementary Table S17). Thus, SQDG falls while PG, the anionic thylakoid lipid it would normally replace, remains flat, and the two together fall from 3.13% to 1.72% of TIC. Phospholipids hold steady (30.3% to 30.2% of TIC) while glycolipids fall (47.5% to 43.5%), raising the ratio of phospholipids to glycolipids. Additionally, all phospholipids increase relative to SQDG (PC 2.07×, PE 2.41×, PG 2.54×; Supplementary Table S17). PC salvage via deacylation would increase LPC, but LPC does not accumulate (0.87 ×CLR), so we find no evidence for that route behind the modest PC decline. Overall, the lipidome suggests a smaller plastid lipid fraction with a disproportionate loss of SQDG. Because SQDG is the only sulfur-containing membrane lipid, constrained sulfolipid synthesis is one explanation for why it falls further than galactolipids; though we did not measure sulfur, and other explanations remain open. The LIN GWAS returned one gene annotated as a sulfoquinovosyl transferase SQD2 (`SORBI_3002G000600`) and one paralogue from the same glycosyltransferase family (`SORBI_3003G076900`), both under fatty-acid traits rather than any SQDG trait, so they are candidates to follow up rather than evidence for the sulfolipid route. Without sulfur, chlorophyll, or thylakoid measurements, the lipidome alone cannot separate these possibilities. Therefore, we describe the LIN lipidome as a redistribution among plastid, phospholipid, lysophospholipid, and neutral-storage pools rather than a phosphate-driven remodeling.

#### Genetic control of lipid variation is stronger under LIN and largely trial-specific

Heritability was low in both trials. It is a ratio, so a small value can indicate little genetic variance or a large residual, and with no replication, we have no independent estimate of plot error to distinguish between the two. Our estimates also come from a marker relationship matrix, so they capture only the additive variance that markers tag and will remain below broad-sense estimates from replicated designs. Published leaf metabolomes provide some backup. Mean broad-sense heritability across 76 wheat leaf metabolites was 0.25, with genotype-by-location variance twice the genetic variance (Matros et al. 2017), and 25–40% of maize leaf mass features fell below *H*^2^ = 0.2 (Zhou et al.2019). Our LIN values sit near that range, and our CTL values are below it, which the narrow-sense comparison explains in part but not fully.

The heritability difference between the trials remains consistent. The paired estimates use the same genotypes and the same species, so it is not an artifact of the panel or the species set. It is unlikely to be a power problem since the genotyped sets are nearly the same size in the two trials, or a noise problem since total phenotypic variation was similar. The data show a smaller genetic share of variance under CTL. Without replication, we cannot tell whether this reflects less genetic variance, more residual variance, or both. This is consistent with the idea that favorable conditions mask genetic differences in lipid composition, which emerge under stress. We did not test which LIN factor drove the effect. Ancestry explains little in either trial, and the small detectable signal appears only under LIN.

The GWAS points the same way. Under LIN, species with candidate genes had a higher median *h*^2^ than those without (0.345 compared to 0.129), while under CTL, both groups were near zero. More heritable traits are easier to map, so this is a consistency check, not independent support. Likewise, the longer LIN candidate list may reflect more mappable traits, not more genes. After collapsing linkage, candidate loci are mostly trial-specific, and repeated regions do not have the same trait.

#### Control-associated loci point to cuticular and phospholipid-turnover genes

CTL returned fewer candidates than LIN, and all 20 well-supported gene sets come from TG traits, with most relating to general cellular processes. Four name a lipid process, and one of those, *triacylglycerol lipase activity*, is treated above with the reaction network. The remaining three fall into two groups, cutin biosynthesis and phospholipid turnover (Supplementary Table S2).

*Cutin biosynthetic process* is the most enriched of the twenty and is driven by `SORBI_3001G222700` and `SORBI_3003G360700`. `SORBI_3001G222700` encodes a very-long-chain aldehyde decarbonylase related to rice WDA1, a wax biosynthesis protein (Jung *et al*. 2006) and associates with eight TG species, five palmitatecontaining. `SORBI_3003G360700` is a GPAT6-family acyltransferase. In tomato, it esterifies hydroxy fatty acids to glycerol, and its loss reduces cutin (Petit *et al*. 2016). Because cutin and wax monomers derive from C16/C18 pools, the palmitate enrichment is consistent with shared precursors. As we measured whole-leaf lipids (not surface wax), this likely reflects precursor pools rather than cuticle composition.

The second group is phospholipid turnover. *Phospholipid dephosphorylation* and *phospholipid metabolic process* share LPP-family phosphatidic acid phosphatases `SORBI_3003G078700` and `SORBI_3010G193300`. The latter term also includes secretory phospholipase A2 `SORBI_3005G145100`. LPPs convert PA to DG, limiting PA signaling. In Arabidopsis, *LPP2* loss elevates PA and increases ABA-sensitive germination (Katagiri *et al*. 2005). Sorghum’s drought tolerance is attributed in part to leaf wax load (Busta *et al*. 2021; Fontanet-Manzaneque *et al*. 2025), and both the cuticle and ABA signaling are under lipid control. Thus, these five genes are worth testing under water deficit. That is a hypothesis rather than a result, since CTL received standard management with sufficient irrigation.

#### Fatty-acid traits under LIN map to a chromosome 3 block of cold- and development-linked regulators

One LIN locus is notable for showing consistent, fatty-acid– specific associations. `SORBI_3003G406800`, a MADS-box transcription factor on chromosome 3, is associated with FA(22:1) and PC(18:2_20:4) and with the fatty-acid class sum plus eight FA-to-class ratios (Supplementary Table S23). The signal does not resolve to one gene, eight genes in a 340 kb window share the same association pattern, and `SORBI_3003G406600` (SBP/SPL) is in perfect LD (*r*^2^ = 1) with the lead SNP. The only LD-aware GO term is *DNA binding* (from 1,166 genes), so interpretation relies on the associations and prior studies.

Three genes in the region have relevant literature. The sorghum MADS gene is cold-induced and a hub in an early shock co-expression module (Kurt 2026). Its closest maize homolog, *ZmMADS69*, is a flowering-time locus involved in tropical-to-temperate adaptation (Liang *et al*. 2019). Additionally, the maize ortholog of the SBP/SPL gene, *ZmSPL6*, accelerates female flowering when knocked out under long days but not short days, implying photoperiod- and temperature-dependent effects (Jin *et al*. 2023). Also in the window, `SORBI_3003G407200` (*r*^2^ = 0.50) encodes a CG-1 (CAMTA) DNA-binding domain. In Arabidopsis, CAMTAs regulate *CBF2* and contribute to freezing tolerance (Doherty et al. 2009).

LIN was planted earlier to increase cold exposure, so a window containing two cold-linked regulators associated with fatty-acid traits is consistent with cold acclimation, which remodels acyl chains to maintain membrane fluidity (Tan et al. 2018; Zhang et al. 2020). However, earlier planting also changes photoperiod and temperature during development, and the *ZmSPL6* phenotype is conditional on such developmental timing, making that explanation equally plausible. However, our design cannot distinguish these effects singularly because of compounding effects. The data support only an association between this block and fatty-acid traits under LIN, and the causal pathway remains unresolved.

#### Nitrate and NPF transporters link nitrogen status to fatty-acid traits under LIN

Nitrogen transport enters the LIN candidate set via two GO enrichments. First, nitrate-specific terms (*nitrate transmembrane transporter activity* and *nitrate transmembrane transport*) are strongly enriched in the fatty-acid set because three of the five annotated genes are candidates. However, all three cluster on chromosome 4. Second, broad transport terms (*transmembrane transporter activity* and *transmembrane transport*) have large backgrounds (402 and 865 genes) and weak enrichment (1.2–2.6×), providing no cargo specificity, but their genes span up to 39 intervals and recur across the fatty-acid, PC, TG, and SQDG sets, implying a dispersed, multi-locus signal.

Nitrate-associated terms map to a 250 kb region on chromosome 4 containing three NRT2-family genes, including `SORBI_3004G009200`, `SORBI_3004G009400`, and `SORBI_3004G009500`. All share the closest maize homolog *nrt2* (`Zm00001eb209670`) and correspond to rice *OsNRT2*.*1*/*OsNRT2*.*2*, high-affinity root nitrate transporters that function with NAR2.1 (Yan et al. 2011; Lupini et al. 2016). `SORBI_3004G009400` associates with FA(22:1), the fatty acid class sum, and the oxidized fatty acid 13S-HOTrE. The other two peak at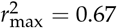 and 0.55, and each associates with one lipid. Because the interval includes 23 LIN candidate genes, LD does not uniquely implicate the transporters.

Two additional NPF/NRT1 transporters, `SORBI_3008G081400` and `SORBI_3001G133900`, appear in the broad transport terms. `SORBI_3008G081400` is recovered in both trials (seven TGs under CTL, five of which are palmitate-containing; FA(22:1), TG(18:1_18:2_18:2) and 11,14,17-eicosatrienoic acid under LIN). Its closest Arabidopsis homolog is NRT1.2, which transports both nitrate and abscisic acid without the two competing as substrates (Kanno et al. 2013). The same protein interacts with PHOSPHOLIPASE D*α*1 at the plasma membrane, with *PLDα1* epistatic to *NRT1*.*2* in ABA responses (Li et al. 2020b), placing the homolog at a junction of nitrate transport, ABA transport and phospholipase D activity, a plausible point of contact between N status and membrane lipid turnover, however, that interaction was characterized in seed germination rather than in leaves. `SORBI_3001G133900` associates with one lipid, TG(14:0_18:2_18:3), and its rice ortholog OsNPF2.4 mediates low-affinity nitrate root-to-shoot transfer (Xia *et al*. 2015). Thus, these transporters are the most promising nitrogen-related candidates in the LIN set for follow-up work.

#### Phosphate-starvation genes and a plastid sulfate transporter associate with fatty-acid and TG traits under LIN

Under LIN, the term cellular response to phosphate starvation is enriched across most lipid sums, and in the SQDG traits, its three genes are `SORBI_3001G028500` on chromosome 1, and `SORBI_3003G088800` and `SORBI_3003G088900` on chromosome 3. `SORBI_3001G028500` encodes an SPX-domain phosphate-status signaling protein and is associated with FA(22:1), the fatty-acid class sum, and the SQDG-to-fatty-acid ratio. Like rice OsSPX4, which negatively regulates phosphate signaling through its interaction with PHR2, it may regulate phosphate-starvation responses (Lv *et al*. 2014).

The two chromosome 3 genes sit in one of the most recurrent loci recovered under LIN, across 52 significant SNPs associated with FA(22:1), the free fatty acid class sum, and nine class-ratio traits. `SORBI_3003G088800` and `SORBI_3003G088900` are separated by 1.8 kb and are closest to the rice *OsLPR2* and *OsLPR1*, the multicopper oxidases of the LOW PHOSPHATE ROOT family, which control the root meristem response to phosphate deficiency. In rice, the family has expanded to five genes within 65 kb, of which *OsLPR3* and *OsLPR5* are induced by Pi deficiency (Cao *et al*. 2016), making the sorghum pair closest to the non-induced members. The same block carries a PHO2-family ubiquitin-conjugating E2, `SORBI_3003G089000`. PHO2/UBC24 is a negative regulator that limits phosphate uptake and root-to-shoot transport and is miR399-cleaved under phosphate deficiency (Aung *et al*. 2006; Huang *et al*. 2013). In Arabidopsis, the RING-type E3 ligase NITROGEN LIMITATION ADAPTATION partners with UBC24 to ubiquitinate PT2 (Park *et al*. 2014), a link that matches our design because LIN reduced both N and P. Under the strictest LD setting (*r*^*2*^≥ 0.9) (Fig. 6C), the block resolves to five genes, each carrying a variant in perfect linkage with the lead SNP.

Two transporters appear only under broad transport terms, not phosphate- or sulfur-specific ones (Supplementary Tables S3 and S4). `SORBI_3006G244000`, a STAS-domain protein assigned to the SULTR3 family, is associated with TG(14:0_18:3_18:3), TG(12:0_18:2_18:3), and PC(18:2_20:4). In Arabidopsis, SULTR3 proteins localize to the chloroplast envelope and mediate chloroplast sulfate uptake (Chen *et al*. 2019). Because SQDG synthesis depends on plastid sulfate import and reduction, SULTR3 may act upstream of the SQD2-family loci noted above in the SQDG pathway. Furthermore, `SORBI_3002G116100` encodes a PHT1-family H^+^/Pi cotransporter. Its closest characterized rice homolog is OsPT4 (*OsPht1;4*), which affects phosphate accumulation and embryo phosphate mobilization (Ye *et al*. 2015; Zhang *et al*. 2015). However, its five associated traits are four TGs and one PC, arguing against a simple phospholipid-to-glycolipid replacement under phosphate limitation. Thus, these loci are prioritized for functional validation for P and sulfur related phenotypes.

#### Jasmonate and oxylipin genes are LIN-specific candidates

Under biotic stress, the dominant lipid signals are oxylipins made from unsaturated fatty acids, and jasmonate, in particular, mediates defense against pathogens (Kuźniak and Gajewska 2024b). This pathway starts with lipoxygenases. In the 13-LOX branch, 18:3 is oxygenated and, after three steps, converted to jasmonate, which is then conjugated to isoleucine to form JA-Ile. JA-Ile binds the COI1–JAZ complex, promoting JAZ repressor degradation and activating defense genes (Li *et al*. 2022). LIN enrichments recover this pathway. Seven candidate genes fall under LIN and none under CTL. Five are lipoxygenases across four loci, including `SORBI_3001G125800` and `SORBI_3001G483400` (chr1), `SORBI_3003G385500` and `SORBI_3003G385900` (26 kb apart; chr3), and `SORBI_3008G191000` (chr8). `SORBI_3001G483400` or-thologs Arabidopsis LOX3, whereas `SORBI_3008G191000` corresponds to LOX1 (9-LOX), spanning both branches. Three other candidates lack characterized orthologs. `SORBI_3009G249900` is orthologous to rice *OsJAR1*, which produces JA-Ile. The *osjar1* mutants show light-dependent coleoptile elongation, indicating developmental as well as defensive roles (Riemann *et al*. 2008). `SORBI_3010G130400` is annotated as a jasmonic acid carboxyl methyltransferase. The rice ortholog *OsJMT1* methylates jasmonate, is induced by brown planthopper, and its overexpression increases methyl jasmonate but reduces herbivore-induced JA/JA-Ile and decreases height and yield, consistent with a growth–defense trade-off (Qi et al. 2016).

Given sorghum’s common fungal disease pressure (e.g., anthracnose, grain mold, charcoal rot, downy mildew, rust) (Khaskheli et al. 2025), an untreated trial could possibly shift this pathway. However, we did not test for any insect damage or disease trials in LIN, so this remains a hypothesis for future work.

## Conclusion

SoLD profiled 266 annotated lipid species in the SAP across two contrasting fields. The lipidomes were broadly similar but differed in four population-wide signatures: SQDG depletion, TG enrichment, a PS-centered phospholipid redistribution, and an LPC/LPE shift toward LPE. Botanical race explained none of these, while genome-wide structure explained only the PS shift. Genomic heritability was higher under LIN (median 0.171) than CTL (median 0), and more of the lipidome could be mapped under LIN, though without replication we cannot distinguish higher genetic variance from lower residual variance.

Because GWAS produced too many candidates to interpret, we prioritized hits by (i) gene sets surviving an LD-aware null in at least two independent intervals, (ii) genes on LINEX reaction branches, and (iii) loci with the strongest signals and most associated lipid traits. The resulting candidates line up with each regime: (i) phosphate signals included a chromosome 3 block with two LOW PHOSPHATE ROOT homologues and a PHO2-family E2, plus a chromosome 1 SPX-domain gene; (ii) N mapped to high-affinity nitrate transporters on chromosome 4; (iii) earlier planting to a chromosome 3 block supported by cold-response and flowering-time evidence; and (iv) the absence of pesticide to jasmonate/oxylipin genes. DG–TG cycling appeared on both sides via *DGAT1* and an SDP1-like patatin lipase. The CTL set was narrower, focused on TG traits, and implicated cuticular lipid synthesis and phospholipid turnover, possibly linked to drought response.

Most loci were trial-specific, and intervals often contained multiple genes. The phosphate block covers 15 genes across 233 kb, the nitrate interval includes 23 candidates, and the chromosome 3 cold locus shares associations with seven neighboring genes. Functional validation should therefore target regions (multiplex editing or block deletion/complementation), using single-gene knockouts mainly where linkage is already broken. Because LIN imposed several stresses at once, separating their effects will require same-year factorial experiments that vary one factor at a time.

## Acknowledgments

The work on this paper and Nirwan Tandukar was supported by the U.S. Department of Energy, Office of Science, Biological and Environmental Research program, Early Career Award Number DE-SC0021889. The work (proposal: https://doi.org/10.46936/10.25585/60008134) conducted by the U.S. Department of Energy Joint Genome Institute (https://ror.org/04xm1d337), a DOE Office of Science User Facility, is supported by the Office of Science of the U.S. Department of Energy, operated under Contract No. DE-AC02-05CH11231. This work was supported by the Research Capacity Fund (HATCH), project award no. 7005660, from the U.S. Department of Agriculture’s National Institute of Food and Agriculture. This work was supported in part by the Science and Technologies for Phosphorus Sustainability (STEPS) Center, a National Science Foundation Science and Technology Center (CBET-2019435). We thank members of the Rubén Rellán Álvarez labs for their contribution to field work and sampling that made possible the research in this manuscript.

