## Supplementary Files for "The Sorghum Lipid Database (SoLD): population-scale lipidomics linking environmental and genetic variation in the Sorghum Association Panel"

**Supplement**

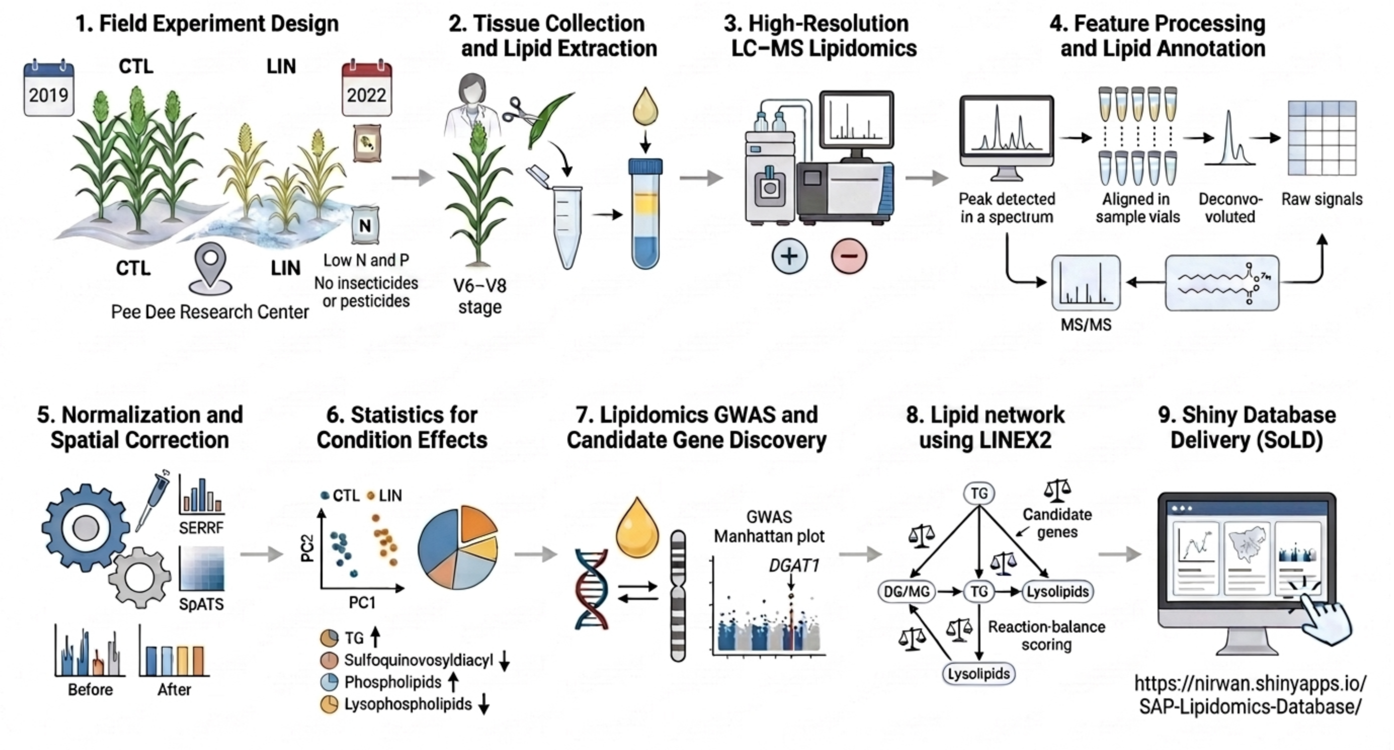


**S1 Fig Field-to-database workflow.** Sample collection and lipid extraction, high-resolution LC-MS lipidomic profiling, GNPS-assisted feature processing and annotation, post-acquisition correction with SERRF and SpATS, statistical description of each trial, lipidomics GWAS and candidate gene discovery, LINEX2 reaction mapping, and dissemination through the SoLD Shiny application.

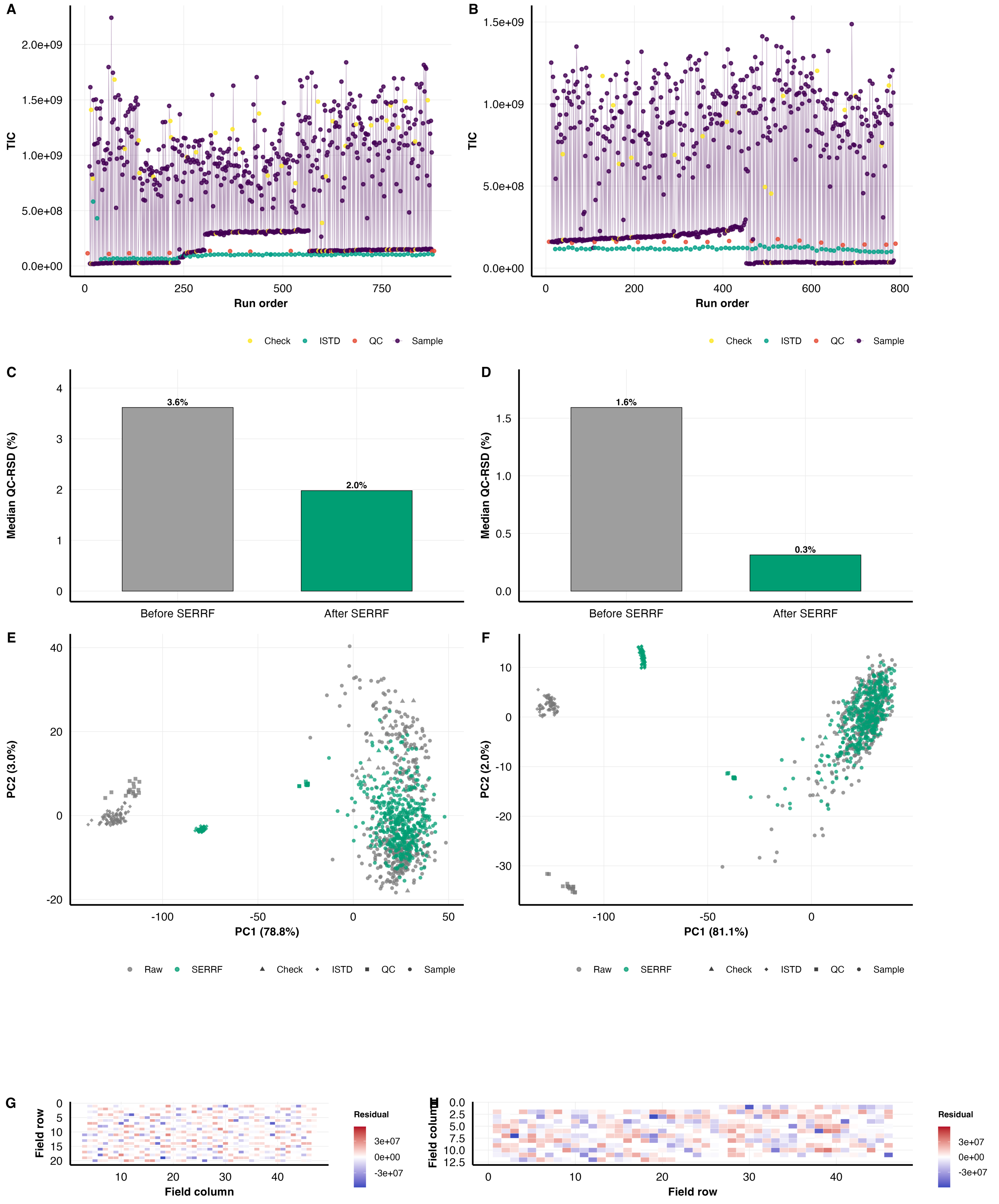


**S2 Fig Quality control, SERRF normalization and SpATS spatial adjustment.** **(A,B)** Run-order TIC for CTL and LIN. **(C,D)** QC-RSD distributions before and after SERRF normalization. **(E,F)** PCA scores before and after normalization. **(G,H)** SpATS spatial residual maps for each trial.

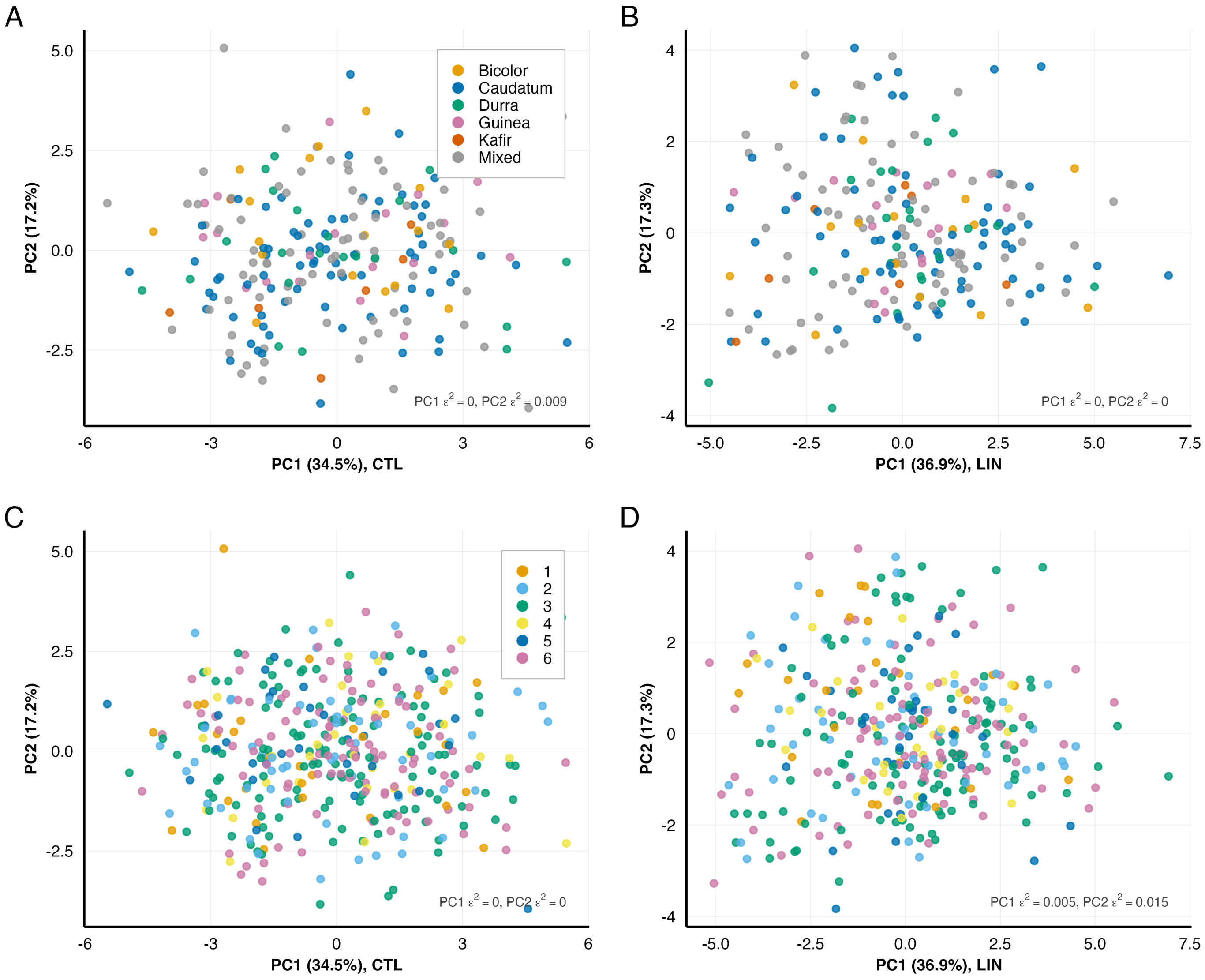


**S3 Fig Lipid-class composition does not separate accessions by botanical race or by genetic cluster.** Principal component analysis of the 13 lipid-class percentages, run within each trial on $\log_{10}$ values scaled to unit variance. **(A,B)** Coloured by botanical race, CTL and LIN. **(C,D)** The same ordinations coloured by marker-based genetic cluster ($K = 6$). Each panel gives the variance in PC1 and PC2 explained by that grouping ($\varepsilon^2$, Kruskal–Wallis).

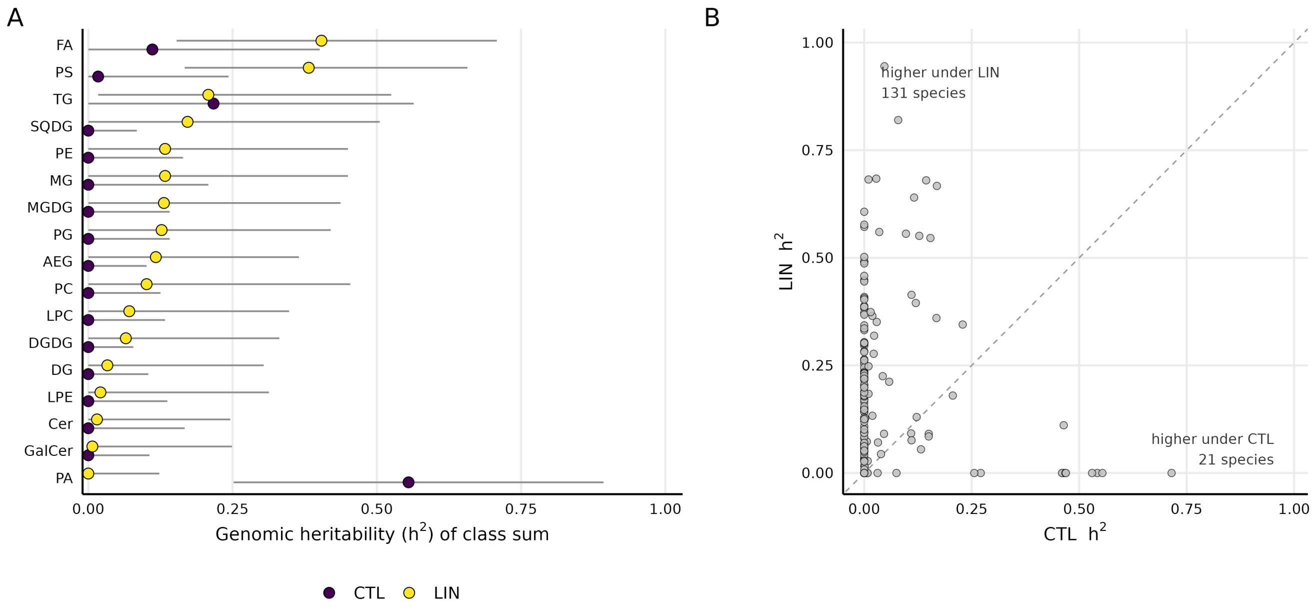


**S4 Fig Genomic heritability of lipid traits in the two field trials. (A)** The 17 lipid-class sums, point estimate with its 95% profile-likelihood interval, on every accession of the trial that is in the relationship matrix. **(B)** Paired per-species heritability, CTL on the horizontal axis and LIN on the vertical, dashed line equality, restricted to the 351 genotypes and 164 species common to both trials. Both panels use the same relationship matrix and the same $\log_{10}$ scale.

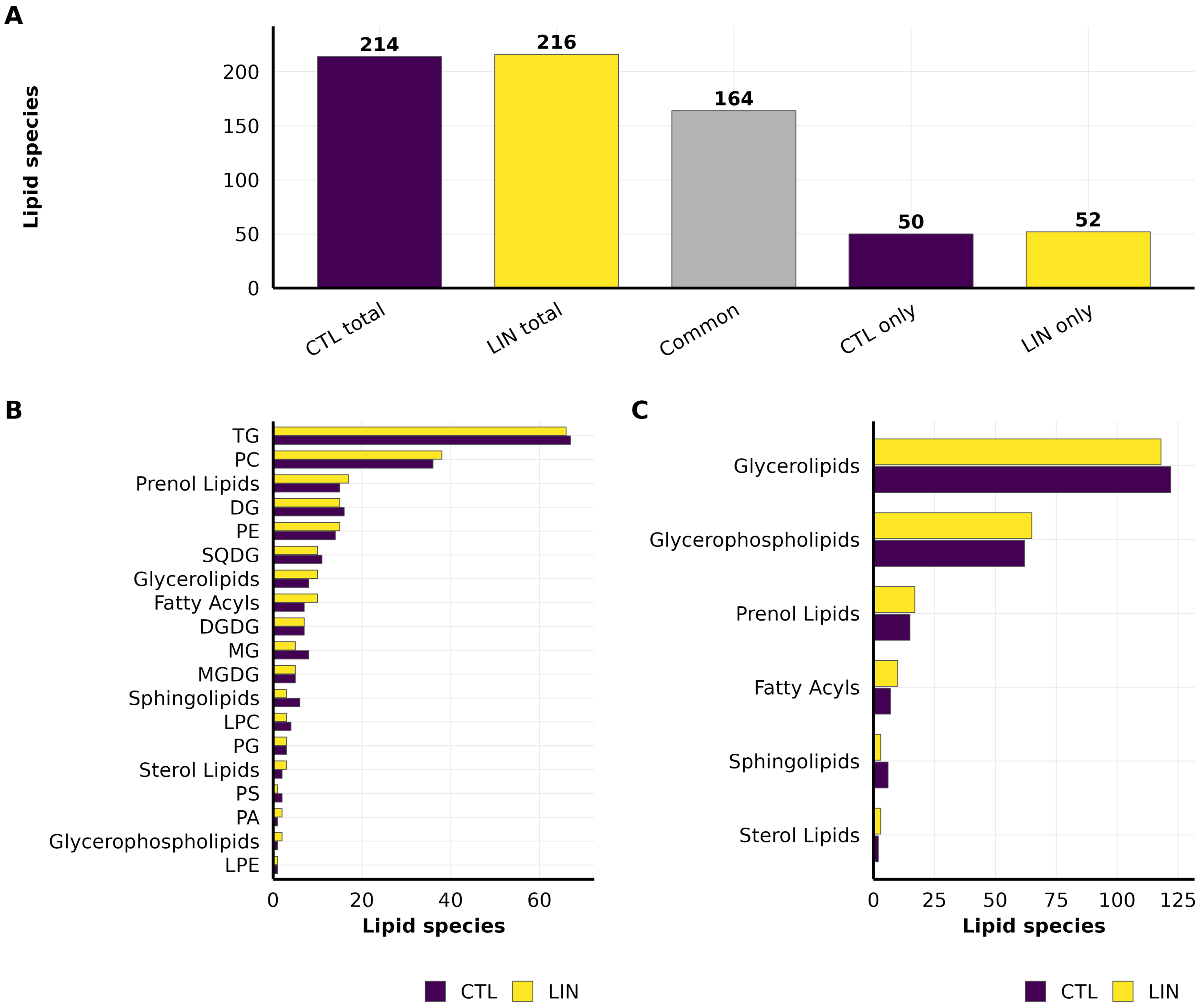


**S5 Fig Lipid species counts and overlap between trials. (A)** Species detected in each trial, with the common and trial-specific counts. **(B)** Counts by lipid class, species with no shorthand name shown under their LIPID MAPS category. **(C)** Counts by LIPID MAPS category. B and C cover every annotated species, so each sums to the totals in (A).

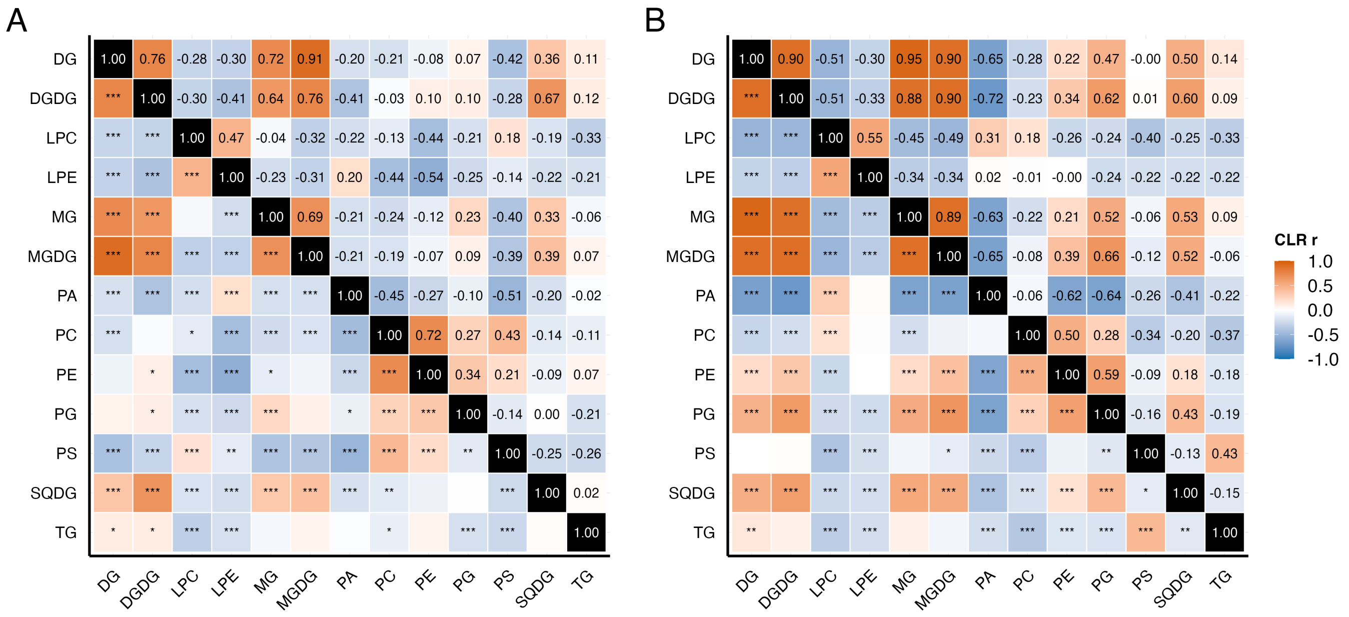


**S6 Fig Correlation structure among lipid classes within each trial.** Pearson correlations on class-level centered log-ratio abundances, **(A)** CTL and **(B)** LIN. Lower triangle, the coefficient; upper triangle, its significance (*$p < 0.05$, **$p < 0.01$, ***$p < 0.001$). Each matrix is computed within one trial; the two are not differenced.

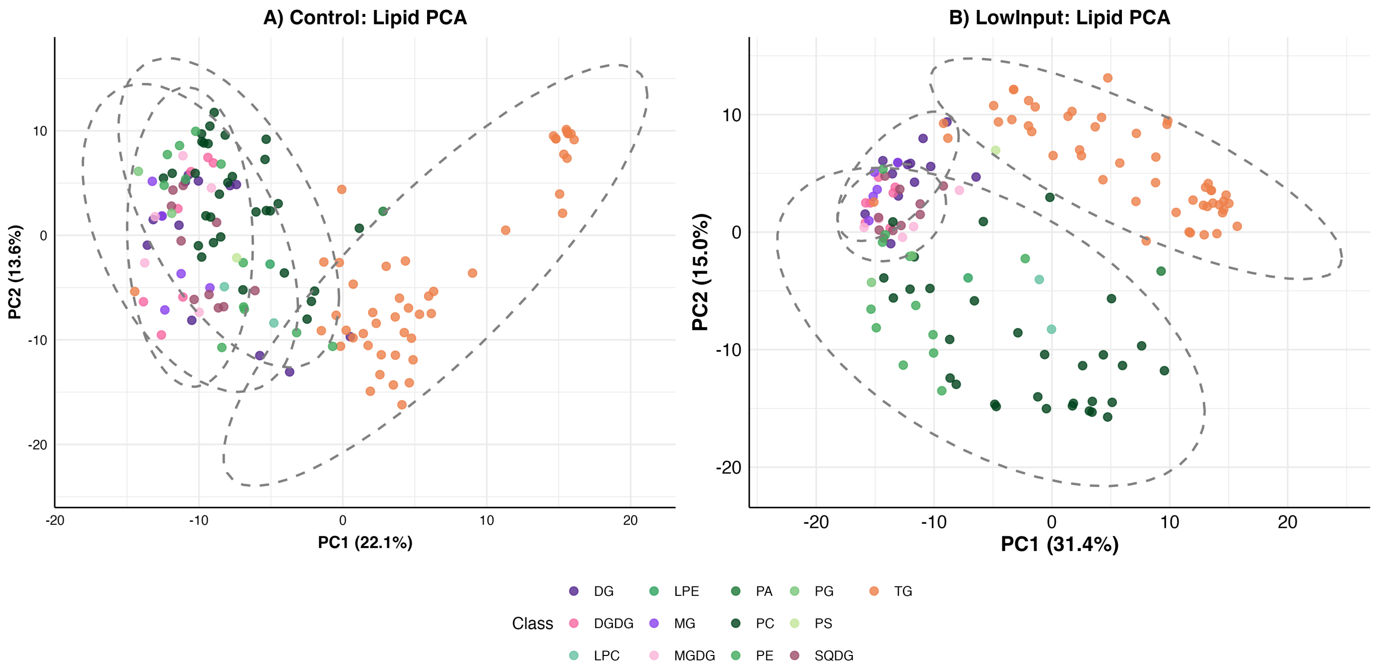


**S7 Fig PCA of lipid species in compositional space. (A)** CLR-transformed lipid species in CTL, with 95% confidence ellipses around four groupings, phospholipids (PC, PE, PG, PA, PS, LPC, LPE), glycolipids (MGDG, DGDG, SQDG), neutral lipids (DG, MG) and storage lipids (TG). **(B)** The same ordination for LIN. Points are coloured by lipid class.

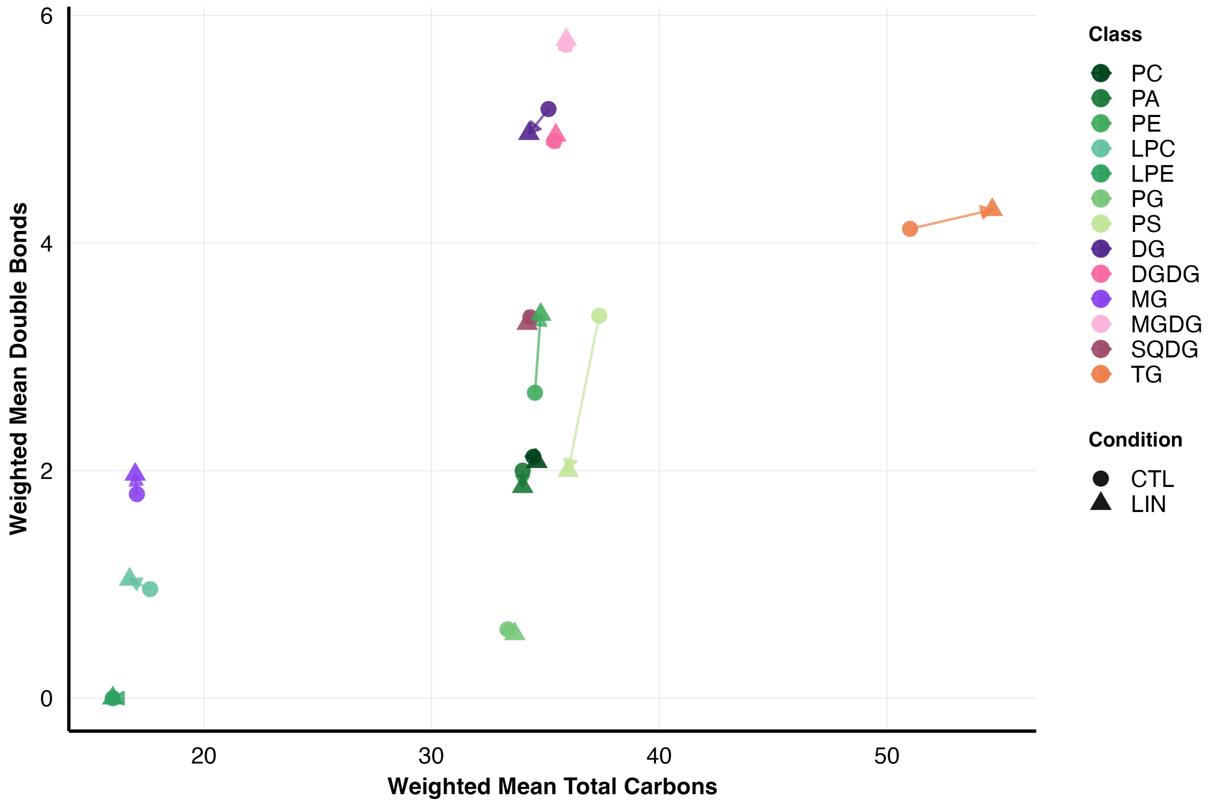


**S8 Fig Lipid classes in a reduced chemical space.** Each point is one lipid class, at its %TIC-weighted mean total carbon number and mean double-bond count across the species it contains. Colour gives the class and shape the trial; the arrow runs from a class's CTL position to its LIN position. The trials differ in year, field and batch, so the shifts are descriptive, and classes represented by one to three species (PS, PA, LPE, PG) have unstable positions because the species detected in the two trials are not the same.

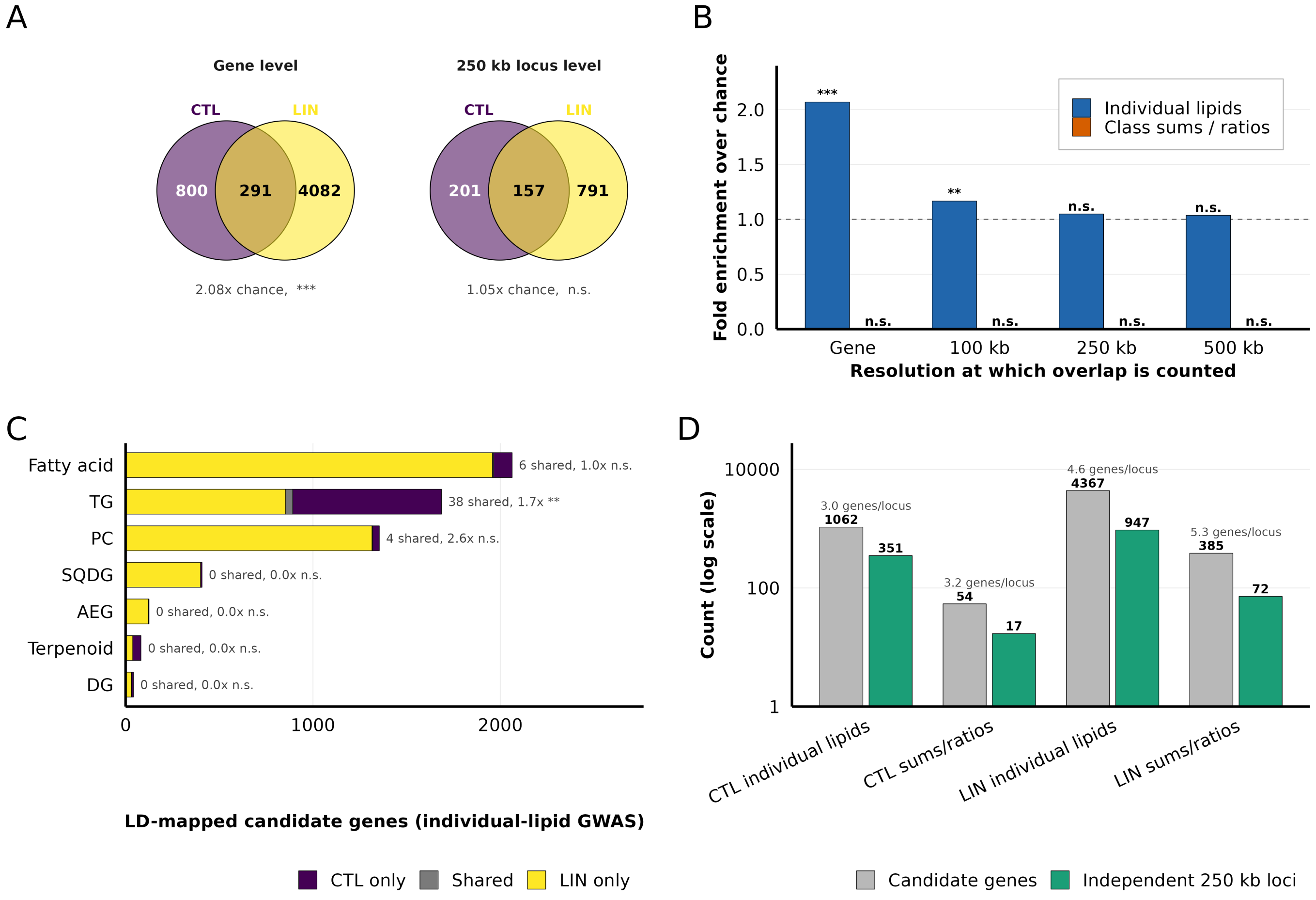


**S9 Fig CTL and LIN GWAS identify largely non-overlapping candidate loci.** **(A)** Overlap of the candidate sets at gene level and after collapsing linkage blocks into 250 kb loci, with fold enrichment and hypergeometric $P$. **(B)** Fold enrichment against the resolution at which the overlap is counted, for individual-lipid and class-sum/ratio traits; dashed line, chance expectation. **(C)** Candidate genes per lipid class in the individual-lipid GWAS, split into CTL-only, shared and LIN-only. **(D)** Candidate-gene counts against the number of independent 250 kb loci they occupy (log scale). *** $P < 0.001$, ** $P < 0.01$, * $P < 0.05$, n.s., not significant.
